# A low-level Cdkn1c/p57^kip2^ expression in spinal progenitors drives the transition from proliferative to neurogenic modes of division

**DOI:** 10.1101/2024.10.10.617342

**Authors:** Baptiste Mida, Nathalie Lehman, Fanny Coulpier, Kamal Bouhali, Rosette Goiame, Morgane Thomas-Chollier, Evelyne Fischer, Xavier Morin

**Author notes:** co-last authors; co-corresponding authors: Electronic address. Electronic address.

## Abstract

During vertebrate neurogenesis, a progressive transition from symmetric proliferative to asymmetric neurogenic divisions is critical to balance growth and differentiation. Using single-cell RNA-seq data from chick embryonic neural tube, we identify the cell cycle regulator Cdkn1c as a key regulator of this transition. While Cdkn1 is classically associated with neuronal cell cycle exit, we show that its expression initiates at low levels in neurogenic progenitors. Functionally targeting the onset of this expression impacts the course of neurogenesis: Cdkn1c knockdown impairs neuron production by favoring proliferative symmetric divisions. Conversely, inducing a low-level CDKN1c misexpression in self-expanding progenitors forces them to prematurely undergo neurogenic divisions. CDKN1c exerts this effect primarily by inhibiting the cyclin D-CDK complex and lengthening G1 phase duration. We propose that Cdkn1c acts as a dual driver of the neurogenic transition whose low level of expression first controls the progressive entry of progenitors into neurogenic modes of division before a higher expression mediates cell cycle exit in daughter cells. This highlights that the precise control of neurogenesis regulators’ expression levels sequentially imparts distinct functions, and is essential for proper neural development.

## INTRODUCTION

The vertebrate central nervous system (CNS) is a complex assembly of thousands of cell types, which are organized in an exquisite manner to form functional neural circuits. This amazing diversity develops through the sequential production of neuronal and glial cells from a limited pool of neuroepithelial stem cells (also called neural progenitors) (Noctor et al., 2001; Taverna et al., 2014). Precise coordination between growth and differentiation during the neurogenic period is paramount to produce the correct number of neural cells “at the right place and time” to ensure the formation of neural circuits. To achieve this complex organization, both the number of times progenitors enter a cell cycle, and the proportion of progenitors that exit the cell cycle to produce neural cells after each round of division are crucial. After initial amplification via proliferative symmetric divisions, the progenitor pool progressively switches to neurogenic modes of division: neurogenic progenitors first perform asymmetric divisions, allowing the self-renewal of one daughter cell while its sibling commits to differentiation, and later switch to terminal symmetric divisions producing two differentiating neural cells (Taverna et al., 2014). Clonal analyses in the mouse and rat embryonic cortices indicate that progenitors that have undergone a neurogenic division do not normally reenter a proliferative state, suggesting an irreversible switch in competence (Gao et al., 2014; Noctor et al., 2004, 2001). This switch was proposed to result from differences in transcriptomic and chromatin landscapes between proliferative and neurogenic progenitors (Aprea et al., 2013; Arai et al., 2011; Haubensak et al., 2004; Iacopetti et al., 1999; Saade et al., 2013; Albert et al., 2017). As an example, expression of the Tis21/Btg2/Pc3 transcription factor (thereafter Tis21) is initiated during the switch from proliferation to neurogenesis from the forebrain (Iacopetti et al., 1999) to the spinal neural tube, in both the mouse (Haubensak et al., 2004) and chick models (Hämmerle et al., 2002; Saade et al., 2013). These observations suggest that self-expanding and neurogenic progenitors correspond to two distinct and successive stages in the neural developmental program.

Here, we generated single-cell transcriptomics data (scRNAseq) from embryonic chick spinal neural tube and identified several genes differentially expressed during the progression from proliferative to neurogenic progenitor states. Our analysis established that Cdkn1c (Cyclin-dependent kinase inhibitor 1c/p57^kip2^) a known regulator of cell cycle exit in newborn neurons (Gui et al., 2007; Mairet-Coello et al., 2012; Tury et al., 2011) is already expressed at low level in neurogenic spinal cord progenitors. Using loss of function and controlled overexpression experiments, we demonstrated that the onset of Cdkn1c expression in cycling progenitors is a driver of the transition towards neurogenic modes of division. Mechanistically, Cdkn1c acts in progenitors by increasing the duration of the G1 phase of cell cycle, mainly via inhibition of the CyclinD1/CDK6 complex. Our findings suggest a dual role for Cdkn1c: a low Cdkn1c expression initially promotes neurogenic modes of division before its higher expression facilitates cell cycle exit in newborn neurons.

## RESULTS

### 1) Transcriptional signature of the neurogenic transition

To identify potential drivers of the transition from the proliferative to the neurogenic state, we produced single-cell transcriptomics (scRNAseq) data from the cervical neural tube region of chick embryos at HH18 (E2.75), when proliferative and neurogenic modes of division are about equally represented (Bonnet et al., 2018; Saade et al., 2013). To overcome limitations in the number of reads assigned to genes resulting from the poor annotation of many genes’ 3’UTRs in the chick genome, we built an improved genome annotation based on the galGal6 reference and bulk long-read RNA-seq from the same tissue (embryonic spinal neural tubes, HH18, see Methods). This considerably improved the assignment of scRNAseq reads to genes and therefore the reliability of expressed genes counts in each cell.

We restricted our analyses from the original dataset to central nervous system-related cells (1878 cells) (Figure 1A), excluding neural crest and mesoderm derivatives (see Methods). In UMAP representations (Figure 1B), neural cells do not arrange in isolated clusters. We therefore defined a scoring system based on the levels of expression of a list of progenitor and neuron-specific genes in each cell (see Methods). This system shows the highest progenitor (P) and neuron (N) scores at opposite ends of the UMAP representation, with intermediate values of both scores in between (Figure 1B), indicating that the progression from progenitor to neuron spontaneously emerges as a strong discriminating factor. Importantly, the expression of Tis21 peaks in the region containing cells with intermediate values of the progenitor score and low neuron score (Figure 1B). Hence, this region likely hosts neurogenic progenitors, in agreement with clonal analyses showing that progenitors undergo a progressive maturation from proliferative to neurogenic states. We refined this analysis through pseudo-temporal classification of these cells to identify gene clusters with similar transcriptomic profiles along the pseudo-time axis (Figure 1C). Interestingly, the gene cluster that contained *Tis21* also contained genes encoding proteins with known expression and/or functions at the transition from proliferation to differentiation, such as the Notch ligand Dll1, the bHLH transcription factors Hes6, NeuroG1 and NeuroG2, and the coactivator Gadd45g. We used this list of genes to define a “neurogenic progenitor” (PN) score (Figure 1D) and performed a differential expression analysis based on P, N and PN scores (Figure 1E). The top 10 differentially expressed genes between PN and the other two populations contains five of the six genes used to define the PN score. In addition, it includes a) three genes with unknown function in the neurogenic transition (*Zc3h12c, LOC107051857, Bambi*), b) the centrosomal gene *Ninein* (*Nin*), whose differential regulation at that stage has previously been described (Zhang et al., 2016), and c) *Cdkn1c* (Figure 1E).

**Figure 1.**
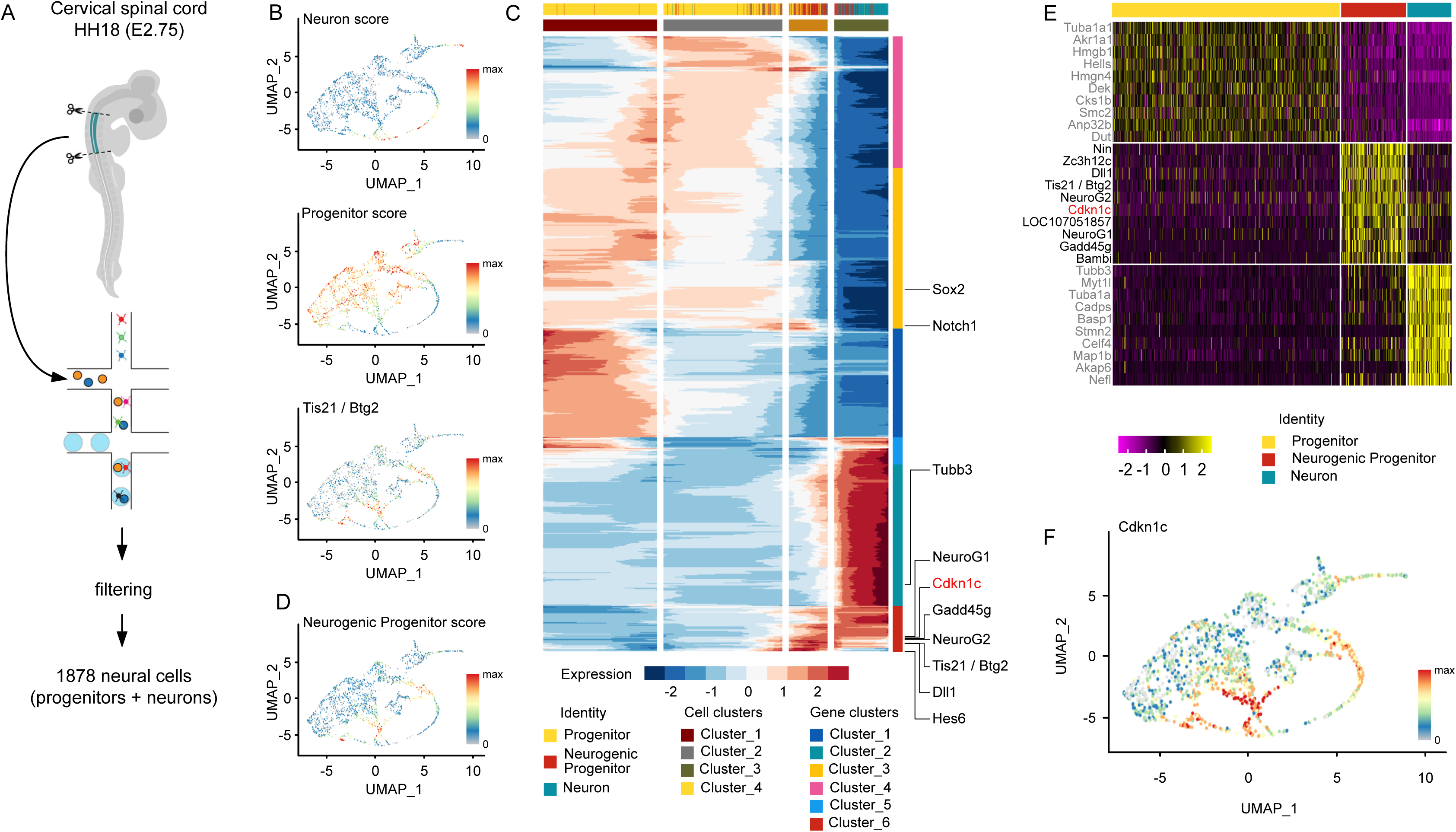
: scRNAseq data analyses from embryonic neural tube led to the identification of Cdkn1c as a potential regulator of the transition of modes of division. **A.** Scheme of the dissection and protocol for scRNA seq generation from chick cervical spinal tube at E2.75. **B.** Visualization of progenitor and neuron scores and of *Tis21/Btg2* expression on the UMAP representation of 1878 chick cervical spinal tube neural cells. **C.** Representation of the pseudotime analysis of the chick scRNAseq dataset. The heatmap shows 6 clusters of genes with similar pattern represented on the vertical axis and 4 cell clusters on the horizontal (pseudotemporal) axis. A subset of genes used to define Progenitor, Neuron, and Neurogenic Progenitor scores are indicated on the right side of the heatmap, illustrating that the three signatures relate to different gene clusters. Cdkn1c is found in the same gene cluster as Neurogenic Progenitor genes. The top horizontal row indicates the cell subtype assigned to each cell along the temporal axis. The blue/red color gradient represents the value of the Z-score. **D.** Visualization of the neurogenic progenitor score on the UMAP representation of 1878 chick cervical spinal tube neural cells. **E.** Heatmap of the 10 most differentially expressed genes between Progenitor, Neurogenic Progenitor and Neuron populations. **F.** Visualization of *Cdkn1c* expression on the UMAP representation of 1878 chick cervical spinal tube neural cells.

Emerging as one of the strong candidates from our analyses, *Cdkn1c* (Cyclin-dependent kinase inhibitor 1c) encodes the p57^kip2^ protein (thereafter named Cdkn1c), and is a classical regulator of G1 phase progression and G1 to S phase transition (Hatada and Mukai, 1995; Matsuoka et al., 1995; Taniguchi et al., 1997). Surprisingly, during CNS development, Cdkn1c has mostly been described as a driver of cell cycle exit (Gui et al., 2007; Tury et al., 2011). Nonetheless, *Cdkn1c* expression has been observed in some neural progenitors (Gui et al., 2007; Mairet-Coello et al., 2012; Tury et al., 2011) and a shortening of G1 phase has been reported in the *Cdkn1c* knock-out mouse cortex (Mairet-Coello et al., 2012). Interestingly, *Cdkn1c* transcripts are specifically enriched in Tis21 positive progenitors in the mouse embryonic cortex (Arai et al., 2011). Accordingly, our pseudotime analyses and UMAP visualization in the chick spinal cord show an onset of its expression in the Tis21 positive neurogenic progenitor population (Figure 1C, F). However, while *Tis21* expression is switched off in neurons, *Cdkn1c* transiently peaks at high levels in nascent neurons before fading off in more mature cells. This peculiar expression profile suggests a dual role of Cdkn1c during neurogenesis: its onset of expression in cycling progenitors would first favor a transition from proliferative to neurogenic modes of division, before higher levels of its expression in daughter cells fated to become neurons would drive them out of the cell cycle.

### 2) Progressive increase in Cdkn1c/p57^kip2^ expression underlie different cellular states in the embryonic spinal neural tube

To test this hypothesis, we explored the dynamics of *Cdkn1c* expression by *in situ* hybridization in the chick embryonic spinal tube (Figure 2A) during the neurogenic transition. While *Cdkn1c* was not expressed at E2, before neurogenesis really starts, its transcript was detected at E3 and E4, when neurogenesis is well underway, as underscored by the expression of the neuronal maker HuC/D in the mantle zone (see lower panels of Figure 2A). It was expressed at low levels in a salt and pepper fashion in the ventricular zone, where the cell bodies of neural progenitors reside, and markedly increased in a domain immediately adjacent to this zone which is enriched in nascent neurons on their way to the mantle zone. In contrast, the transcript was completely excluded from the mantle zone, where HuC/D positive mature neurons accumulate. This is consistent with the dynamics of the *in silico* profile described above.

**Figure 2.**
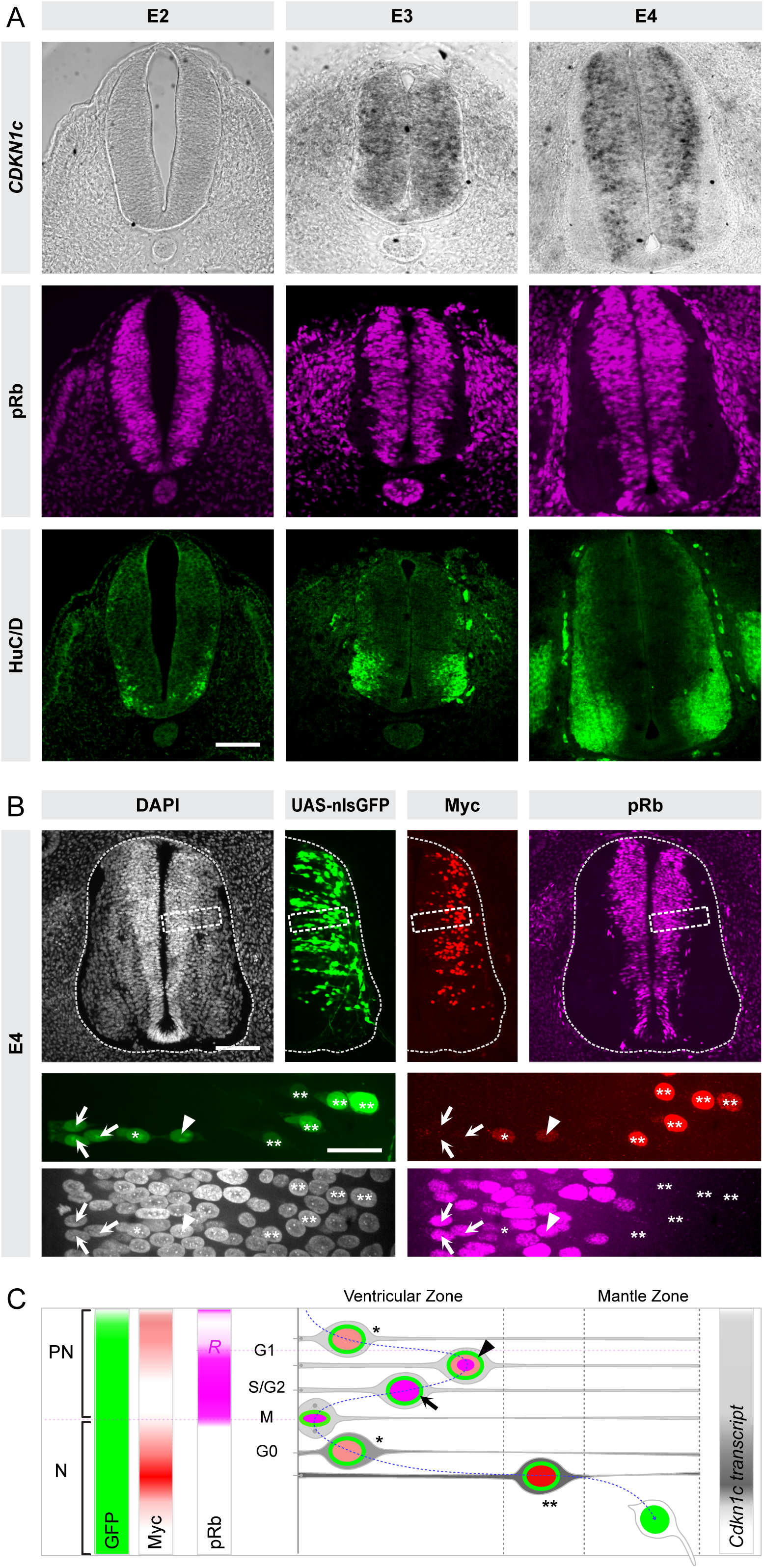
: Dynamics of Cdkn1C expression during spinal tube neurogensesis. **A. mRNA expression of *Cdkn1c* increases during neurogenesis in chick embryonic neural tube**. *In situ* hybridization of Cdkn1c (top row) and immunostainings for pRb (middle row, magenta) and HuC/D (bottom row, green) on cryosections of thoracic region of chick embryonic neural tube at sequential days during development (from left to right: E2, E3, and E4 respectively). For each stage, light and multichannel confocal fluorescence imaging are from a single section. Scale bars: 50µm. **B. Cdkn1c protein is detected at low level in cycling progenitors**. Somatic knock-in of 6xMyc-tags fused to the C-terminus of *Cdkn1c* allows the visualization of Cdkn1c protein (Myc, Red) on E4 transverse vibratome sections. Inclusion of a Gal4-VP16 transcription factor in the knock-in construct (see Supplementary Figure 1) identifies all the cells which express or have previously expressed Cdkn1c via the UAS-nlsGFP reporter (green). Staining with anti phospho-Rb antibody (pRb, magenta) reveals cycling progenitors from late G1 to M phase. The bottom part of the panel shows a close-up of the region highlighted by a dashed rectangle in the top panels. The key to the meaning of asterisks, arrows and arrowheads pointing to cells with different combinations of the markers is illustrated in the scheme in panel C. Scale bars: 50µm and 10µm in close ups. The top row shows maximal projections of 5µm z-stacks; close ups in bottom rows are single confocal z-planes **C. Scheme summarizing the dynamic expression levels of *Cdkn1c* transcript and protein in cycling progenitors and newborn neurons**, as deduced from scRNAseq, *in situ* hybridization and somatic knock-in experiments. In a subset of neurogenic progenitors (PN), the *Cdkn1c* transcript is expressed at low levels (light gray), before it peaks transiently in newborn neurons (N, dark gray) and fades of in more mature neurons (N, white). Cdkn1c protein, visualized with the anti-Myc signal (red) is present at low levels in early G1 in neurogenic progenitors (light red nuclear signal, black asterisk) and shortly overlaps after the restriction point (R) with pRb staining (black arrowhead, light red and magenta nuclear signals). The Myc signal disappears in S/G2 (black arrow) and M phases, during which pRb is still detected (magenta nuclear signal). In newborn neurons, the Cdkn1c /Myc signal is initially detected at low level (light red) and later peaks at its maximal intensity (double asterisks, dark red nucleus) during the early phases of differentiation, before fading out in mature neurons. pRb is absent in the neuronal population. The GFP signal (green) expressed from the UAS reporter is detected throughout this temporal sequence. See main text for details.

We then checked whether the Cdkn1c protein is translated in the population of progenitors in which the transcript is detected. In the absence of functional antibodies in the chick embryo, we used a clustered regularly interspaced short palindromic repeats (CRISPR)/Cas9-based somatic knock-in strategy to insert an array of six Myc tags at the C-terminus of *Cdkn1c* (Supplementary Figure 1A-B and Methods). The Myc tags insertion approach offers a direct read-out of the presence of the protein, and should report any cell cycle dependent stabilization or degradation of Cdkn1c. The Myc tags were immediately followed by a P2A pseudo-cleavage site and the *Gal4-VP16* transcription factor sequence (Supplementary Figure 1A-B) whose simultaneous translation can be used to activate the transcription of a stable fluorescent reporter from a UAS promoter. This allows to identify both Myc-Cdkn1c-positive cells and cells in which Myc-Cdkn1c is no longer present but has previously been expressed.

We electroporated the knock-in vector together with a plasmid expressing Cas9 and guide RNA sequences (gRNAs) targeting the *Cdkn1c* C-terminus and a UAS-nlsGFP (nuclear localization sequence Green Fluorescent Protein) reporter plasmid in the neural tube of E2 chick embryos. At E3, we observed a strong GFP signal in the electroporated side of the embryo using three different *Cdkn1c* gRNAS, whereas no signal was observed when a control gRNA that does not target any chick sequence was used, indicating successful and specific knock-in events (Supplementary Figure 1C). Anti-Myc immunofluorescence on transverse sections at E4 revealed low but detectable signal in the ventricular zone, while a stronger signal was observed in the intermediate domain, and virtually no signal in the mantle zone (Figure 2B). This pattern is very similar to the expression of the transcript observed via *in situ* hybridization (Figure 2A).

To ascertain that Cdkn1c is translated in cycling progenitors, we performed anti-Myc immunofluorescence in combination with an anti-pRb antibody, recognizing a phosphorylated form of the Retinoblastoma (Rb) protein. Although pRb is specific for cycling cells, it is only detected once cells have passed the point of restriction during the G1 phase. We detected many double GFP positive/pRb positive cells (arrows and arrowheads in Figure 2B) in the ventricular zone, indicating that the *Cdkn1c* transcript is translated in cycling progenitors. Only a few GFP positive/pRb positive cells were also Myc positive (arrowheads in Figure 2B). This is consistent with a short period of overlap between Cdkn1c and pRb around the timing of the restriction point, suggesting that Cdkn1c is degraded in later phases of the cell cycle. We also observed some weak Myc positive cells in the ventricular zone that were pRb negative (Figure 2B; asterisks). This may correspond to progenitors in early G1 or alternatively, to early nascent neurons about to upregulate their Cdkn1c expression (see Figure 2C). Finally, we observed GFP positive/pRb negative nuclei with a strong Myc signal in the intermediate domain, corresponding to immature neurons exiting the cell cycle and on their way to differentiation (Figure 2B; double asterisks).

Importantly, these observations confirm that Cdkn1c is expressed at low level in a subset of progenitors in the chick spinal neural tube, as previously suggested (Gui et al., 2007; Mairet-Coello et al., 2012). Our dual Myc and Gal4 knock-in strategy which reveals the history of *Cdkn1c* transcription and translation suggests that the protein is quickly degraded during or after G1 completion. This may explain why a classical immunohistochemistry approach with an anti-Cdkn1c antibody only detected the protein in very few progenitors in the developing mouse CNS (Gui et al., 2007; Mairet-Coello et al., 2012).

Altogether our scRNAseq analyses, *in situ* hybridization and knock-in experiments are consistent with our hypothesis of two sequential roles of *Cdkn1c* in progenitors and neurons.

### 3) Downregulation of Cdkn1c in neural progenitors delays the transition from proliferative to neurogenic modes of division

To functionally investigate the role of Cdkn1c as a potential player in the transition between division modes, we reduced its expression in progenitors. We used a knock-down strategy based on the short hairpin RNA interference (shRNA) approach, with the aim to abolish its low-level expression in neurogenic progenitors, while only partially affecting the higher level of expression required in newborn neurons to trigger cell cycle exit. Of the six shRNA that were tested against *Cdkn1c* (see Methods), only two (shRNA1 and shRNA4) induced an observable reduction of *Cdkn1c* mRNA expression on transverse sections, while an effect of the other four was not clearly visible (Supplementary Figure 2A).

We first investigated the effect of this modest *Cdkn1c* knock-down on the production of neurons at the tissue level. In order to target and investigate specifically the neurogenic transition, we concentrated our analyses on the dorsal region where this transition has barely started at the time of electroporation of the shRNA vectors (E2.25, HH13-14). Neuron and progenitor populations were evaluated 24 or 48 hours after electroporation (hae) via immunofluorescence (see Methods for the choice of the markers of these populations).

shRNA1 and shRNA4 led to a significant increase in the number of pRb positive progenitors 48 hae (Figure 3B and Supplementary Figure 2B). Accordingly, the neuron population, identified with HuC/D staining, was decreased 48 hours after *Cdkn1c* knock-down by shRNA1 (Figure 3B). This switch towards proliferation was already apparent 24 hae, as illustrated by a modest but significant increase in the pRb positive progenitor population in shRNA1 condition (Figure 3A). The reduction in neuronal production was not due to cell death, as we observed no excess in the number of Caspase3 positive cells in the knock-down condition (data not shown). The following experiments were carried out using shRNA1 (thereafter called Cdkn1c shRNA) which showed the most efficient downregulation in *Cdkn1c* expression and the most significant dysregulation in the ratio of progenitor versus neuron populations in our functional studies.

**Figure 3.**
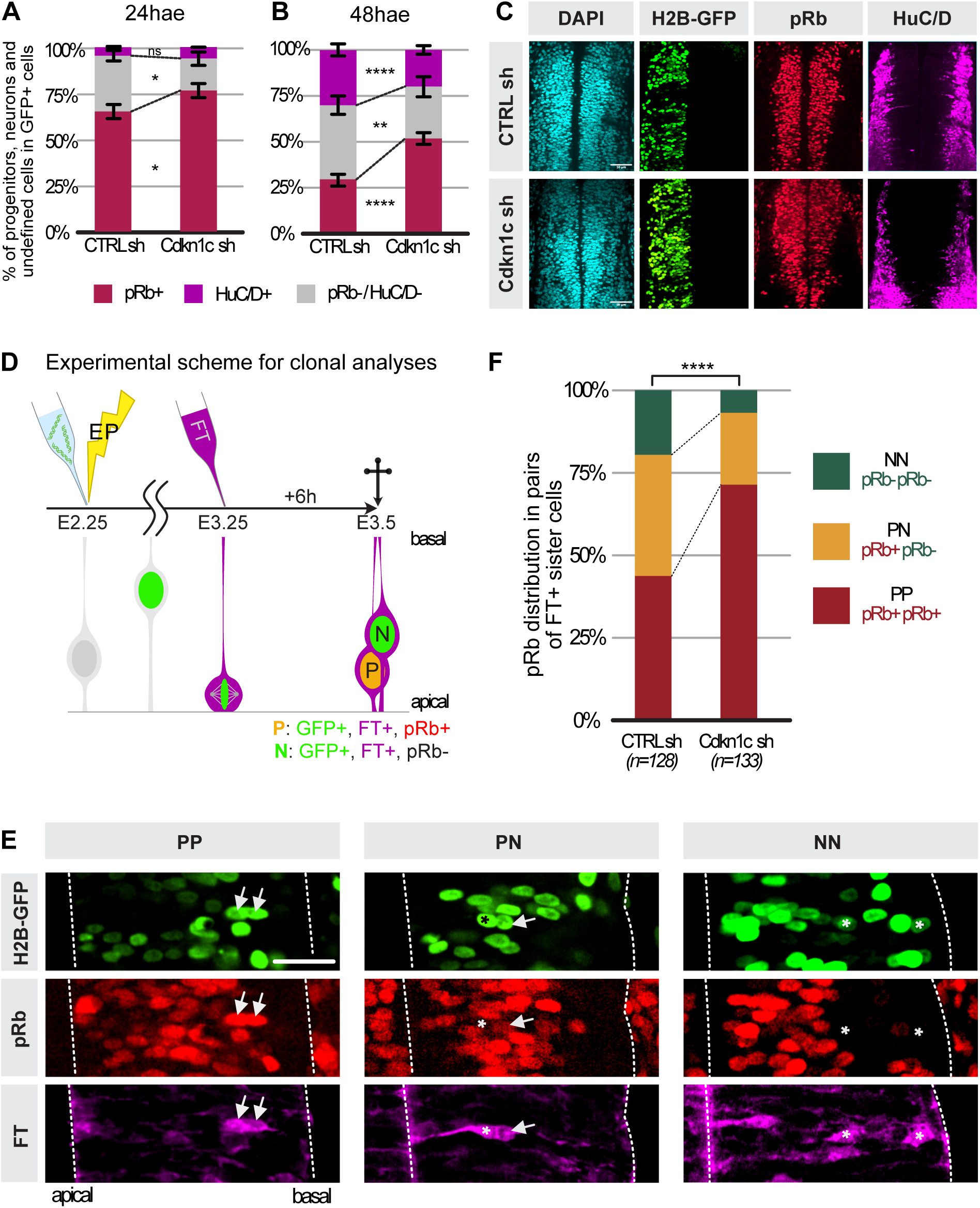
: Downregulation of Cdkn1c delays the neurogenic transition in the spinal cord by favouring a proliferative symmetric mode of division. **A. Distribution of the pRb positive progenitors (dark red), HuC/D positive neurons (magenta) and undefined cells (double pRb/HuC/D negative, gray) in Cdkn1c or control shRNA conditions at E3 24 hours after electroporation (hae).** The undefined population contains both progenitors before the restriction point (and therefore negative for pRb) and immature neurons that do not yet express HuC/D. ns, p > 0.05; *, p < 0.05, (unpaired Student t test). **B. Distribution of the pRb positive progenitors (dark red), HuC/D positive neurons (magenta) and undefined cells (double pRb/HuC/D negative, gray) in Cdkn1c or control shRNA at E4 48 hours after electroporation (hae)**. ns, p > 0.05; **, p < 0.01, ***, p < 0.001 (unpaired Student t test). **C.** Transverse sections of the chick neural tube (thoracic level) at E4 (HH st22) stained with HuC/D antibody (magenta) to label neurons and pRb (red) antibody to label progenitors in **Cdkn1c or control shRNA (sh)** conditions. Scale bar: 50μm. **D. Principle of the analysis of pairs of sister cells.** Embryos are electroporated at HH13-14 (top left, yellow thunder) with Cdkn1c or control shRNA plasmids co-expressing a H2B-GFP reporter. Embryos are injected with the FlashTag dye (FT) 24 hours after electroporation to label a synchronous cohort of mitotic progenitors, and collected six hours later. Anti-pRb and anti-GFP immunofluorescence on thoracic vibratome sections determines the progenitor (pRb positive) or neuron (pRb negative) status of FlashTag-positive electroporated sister cells. **E. Representative examples of two cell clones** in transverse neural tube sections. From left to right panels: PP, PN and NN pairs. Arrows show pRb positive (red) progenitors and asterisks show pRb negative neurons in FlashTag positive (magenta) pairs of GFP-positive (green) sister cells. Scale bars: 25µm **F. Diagram indicating the percentage of PP, PN, and NN clones for control and shRNA electroporated embryos**. PP, PN, and NN stand for divisions producing two progenitors, one progenitor and one neuron, or two neurons, respectively. The distribution of PP, PN and NN clones between control and shRNA was compared using a Chi-2 test, ****p < 0.005.

We postulated that the observed reduction in neurogenesis may result from a delay in the transition from proliferative to neurogenic modes of division, rather than the proposed failure of prospective neurons to exit the cell cycle (Gui et al., 2007). To more directly address this hypothesis, we developed a clonal analysis strategy to analyze the fate of pairs of sister cells born from the division of mother cells downregulated for *Cdkn1c* (Figure 3D). This approach allows to retrospectively deduce the mode of division used by the mother progenitor cell. We injected the cell permeant dye “FlashTag” (FT) at E3 to specifically label a cohort of progenitors that undergoes mitosis synchronously (Baek et al., 2018; Telley et al., 2016 and see Methods), and analyzed the fate of their progeny six hours later (Figure 3D), when two-cell clones selected on the basis of FT incorporation can be categorized as PP, PN, or NN based on pRb positivity (P) or not (N) (see Methods, Figure 3E and Supplementary Figure 3A). Strikingly, upon *Cdkn1c* knock-down, we observed a massive increase in the number of PP clones at the expense of PN and NN clones (Figure 3F). These knock-down experiments comfort the hypothesis that low levels of Cdkn1c favor the transition to neurogenic progenitors. In agreement with this conclusion, analysis of sister cells born from Cdkn1c positive progenitors (identified by knock-in of the Gal4 driver in the *Cdkn1c* locus) showed a significantly higher proportion of neurogenic pairs (PN and NN) compared to the total progenitor population (Supplementary Figure 3B).

### 4) Inducing a premature expression of Cdkn1c in progenitors triggers the transition to neurogenic modes of division

We next explored whether low Cdkn1c activity is sufficient to induce the transition to neurogenic modes of division. To circumvent the cell cycle arrest that is triggered in progenitors by strong overexpression of Cdkn1c (Gui et al., 2007), we developed an approach to prematurely induce in proliferative progenitors the modest level of expression observed in neurogenic progenitors. We took advantage of the *Pax7* locus, which is expressed in progenitors in the dorsal domain at a level similar to that observed for Cdkn1c in neurogenic precursors (Supplementary Figure 4A).

We used the CRISPR/Cas9-based somatic approach to introduce a sequence including a Myc-tagged *Cdkn1c* coding sequence and the *Gal4-VP16* transcription factor downstream of *Pax7* (Figure 4A-B, Supplementary Figure 4B-E). We first confirmed that the Myc signal resulting from the *Cdkn1c-Myc* knock-in at the *Pax7* locus was restricted to progenitors in the dorsal domain. Importantly, its intensity was similar to the one observed for endogenous Myc-tagged *Cdkn1c* in progenitors (Figure 4B and Supplementary Figure 4E), and remained below the endogenous level of Myc-tagged *Cdkn1c* observed in nascent neurons, confirming the validity of our strategy.

**Figure 4.**
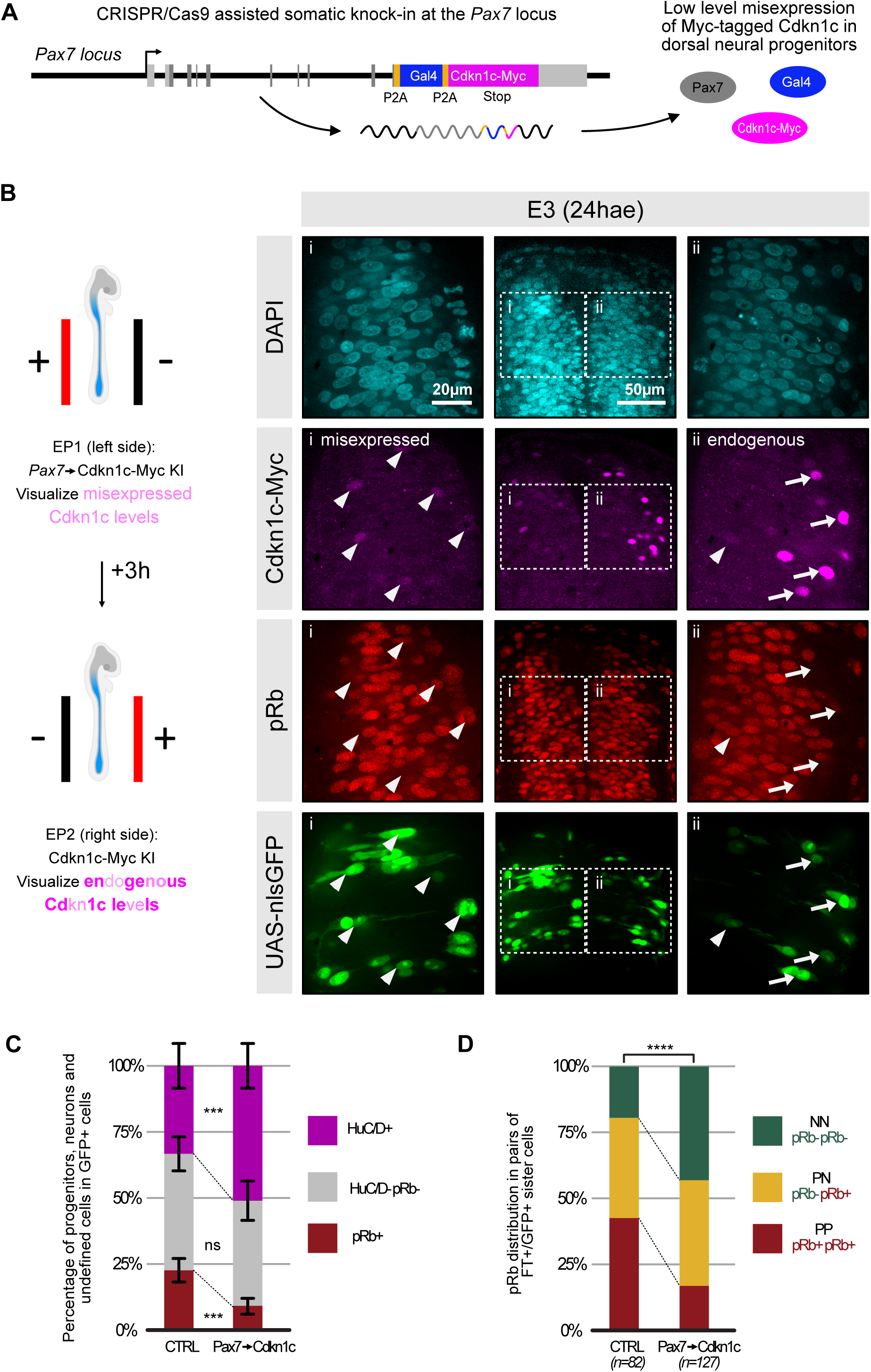
: Premature expression of Cdkn1c at low levels in proliferative progenitors converts them to a neurogenic mode of division **A. Schematic representation of the modified *Pax7* locus driving low-level expression of Cdkn1c-Myc and Gal4-VP16 in dorsal progenitors.** *Pax7* (gray), *Gal4-VP16* (blue) and *Cdkn1c-Myc* (magenta) coding sequences are transcribed from the *Pax7* locus and co-translated. The insertion of P2A pseudo-cleavage sites (orange) between the three sequences ensures that all three proteins are present as independent proteins (see Supplementary Figure 4 and Methods for further details) **B. *Pax7*-driven misexpression of Cdkn1c in dorsal progenitors mimics the levels of endogenous Cdkn1c expression in neurogenic progenitors.** A bilateral electroporation scheme was used to compare *Pax7*-driven levels of Cdkn1c expression (electroporation 1, right side hemi-tube, knock-in of Cdkn1c-Myc in the *Pax7* locus) with endogenous Cdkn1c levels (electroporation 2, left side hemi-tube, knock-in of a Myc tag in the *Cdkn1c* locus). The Cdkn1c-Myc (magenta) signal driven by *Pax7* is restricted to the ventricular region and its intensity is comparable to the endogenous levels of Cdkn1c-Myc in the contralateral side (i and ii, arrowheads) and never reaches the high levels of endogenous Cdkn1c-Myc signal observed in the intermediate zone (i, arrows). Knock-in cells are identified via the expression of a co-electroporated UAS-nlsGFP reporter (green). **C. Distribution of pRb positive progenitors (red), HuC/D positive neurons (green) and undefined cells (double pRb/ HuC/D negative, gray)** in control versus *Pax7*-driven Cdkn1c misexpression conditions 48 hours after electroporation (hae). The control condition consists in a Gal4 knock-in at the *Pax7* locus. For both conditions, knock-in cells were identified via the expression of a co-electroporated UAS-nlsGFP reporter. ns, p > 0.05; *** p < 0.005, unpaired Student t test. **D. Distribution of PP, PN, and NN pairs of sister cells** in control versus *Pax7*-driven Cdkn1c misexpression conditions. The control condition consists in a Gal4 knock-in at the *Pax7* locus. For both conditions, knock-in events in dorsal progenitors were revealed thanks to a co-electroporated UAS-nlsGFP reporter. For the control condition, we used a knock-in construct where Gal4 alone is inserted downstream of *Pax7*. PP, PN, and NN stand for divisions producing two progenitors, one progenitor and one neuron, or two neurons, respectively. The distribution of PP, PN and NN clones between both conditions was compared using a Chi-2 test, ****p < 0.05.

We therefore proceeded to analyze the consequences of Cdkn1c premature expression in progenitors. At the population level, at E4, Cdkn1c expression from the *Pax7* locus resulted in a strong reduction in the number of progenitors (pRb positive cells) and a significant increase in the proportion of neurons (HuC/D positive) within the knock-in population (revealed with a UAS-nlsGFP reporter) (Figure 4C).

We next examined whether the increased neurogenesis 48 hae is linked to a change in the mode of division of progenitors misexpressing Cdkn1c. Using the FlashTag cohort labeling approach described above, we traced the fate of daughter cells born 24 hae. We observed a massive increase in the proportion of neurogenic (PN and NN) divisions rising from 57% to 84% at the expense of proliferative pairs (43% PP pairs in controls versus 16% in misexpressing cells, Figure 4D). Overall, these data show that inducing a premature low-level expression of Cdkn1c in cycling progenitors is sufficient to accelerate the transition towards neurogenic modes of division.

### 5) The proneurogenic activity of Cdkn1c in progenitors is mediated by modulation of cell cycle dynamics

Previous data in the developing cortex of *Cdkn1c* knock-out mice described a transient increase in proliferation (between E14.5 and E16.5) linked to a shorter progenitor cell cycle duration mainly due to a reduction of the G1 phase length (Mairet-Coello et al., 2012). However, in addition to its role as a cell cycle regulator, Cdkn1c also performs other functions, as illustrated by its role as a transcription co-factor or pro-or anti-apoptotic factor in different contexts (Creff and Besson, 2020).

To assess the mechanisms of action of Cdkn1c in the neurogenic transition, we monitored cell cycle parameters in *Cdkn1c* knock-down conditions (see Methods). Cumulative EdU incorporation in spinal progenitors (pRb positive) at E3 (24 hae) showed that the proportion of EdU-positive progenitors reached a plateau faster and in a sharper manner in the shRNA Cdkn1c population (Figure 5A-B). This indicates that the total duration of the cell cycle is shorter upon *Cdkn1c* knock-down. To specifically assess a possible reduction in G1 length, we developed an approach that provides a direct measurement of G1 duration, contrary to the classical method of G1 inference (Nowakowski et al., 1989). Our approach uses precise landmarks to delineate G1 phase borders: mitosis exit (through the FlashTag (FT) labeling of a synchronized cohort of dividing progenitors) and S phase entry (through cumulative EdU labeling; Figure 5C, see Methods). We found that the proportion of pRb positive progenitors having entered S phase (EdU positive cells) was higher at all time points examined after FT injection in the *Cdkn1c* knock-down condition compared to the control population (Figure 5D). Importantly, virtually all progenitors electroporated with Cdkn1c shRNA had reached S phase 10h30 after FT injection, whereas approximatively one-third of control progenitors were still in G1 at that time. Taken together, these results demonstrated a shorter cell cycle in *Cdkn1c*-downregulated condition at the population scale, which is, at least partially, related to a decrease in G1 duration.

**Figure 5.**
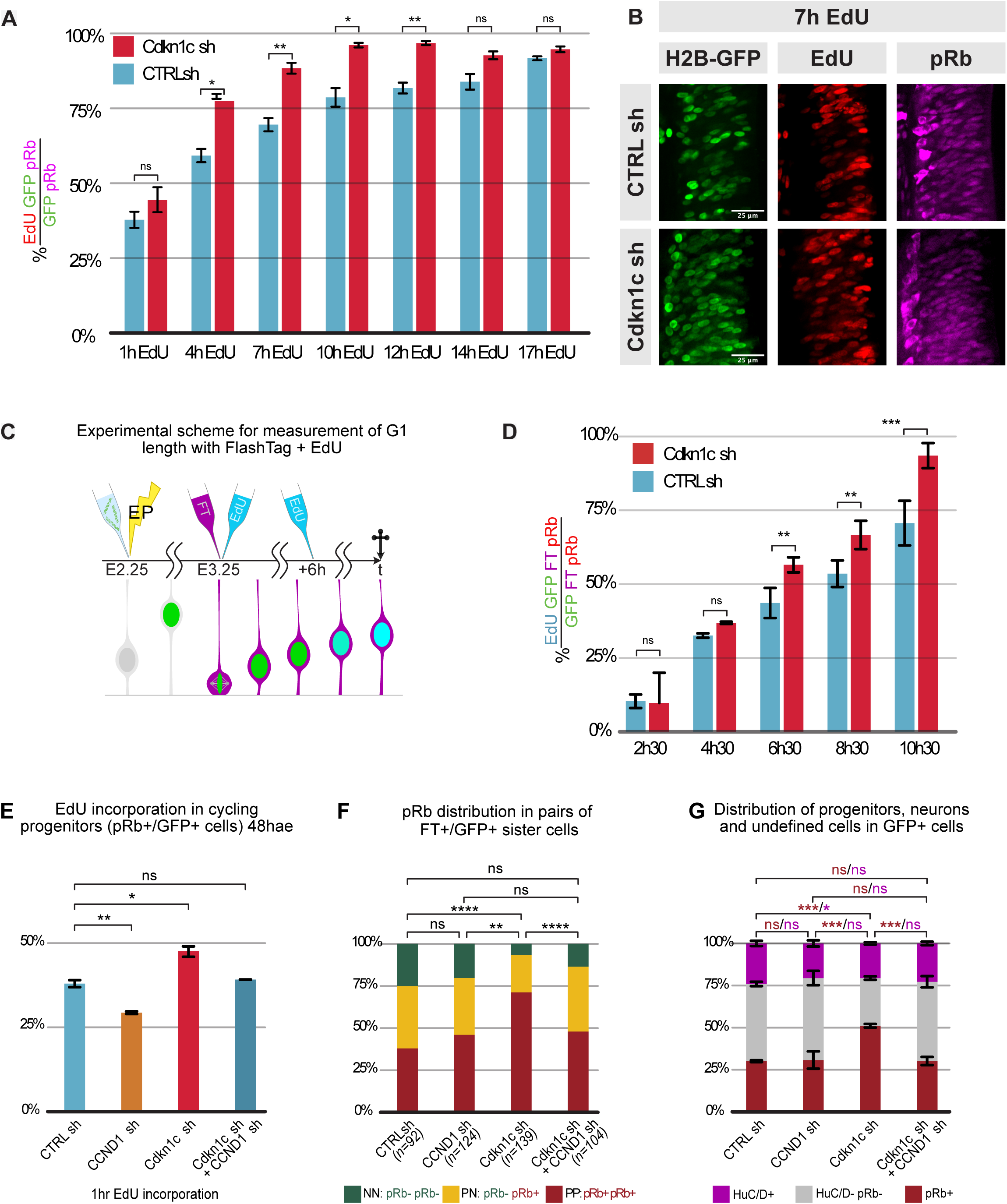
: Cdkn1c controls G1 phase duration in neural progenitors, and acts on cell cycle duration and neurogenesis via regulation of *CCND1* activity. **A. Cumulative EdU incorporation in the overall electroporated progenitor population**. The columns represent the percentages of EdU/pRb/GFP triple positive cells in the electroporated progenitor (GFP/pRb double positive cells) population of control (blue) *versus* Cdkn1c shRNA (red) conditions at each time point. EdU, 5-ethynyl-20-deoxyuridine; ns: p > 0.05; *, p < 0.05; **, p < 0.01 (unpaired Student t test). **B.** Representative images from the EdU incorporation experiment in transverse sections at the 7h timepoint. A close-up of the electroporated side is shown. **C. Schematic representation of the experimental strategy to measure G1 length**. Embryos were electroporated with Cdkn1c or control shRNA at E2.25. One day later, FlashTag injection in the neural tube and EdU administration were performed simultaneously. Embryos were harvested at consecutive time points every 2 hours between 2h30 and 10h30. For time points beyond 6 hours, a second EdU injection was performed 6 hours after the first one. **D. Dynamics of EdU incorporation in a FlashTag-positive cohort of electroporated progenitors.** The columns represent the percentages of EdU-positive cells in FlashTag/pRb/GFP triple positive cells in Cdkn1c (red) and control shRNA (blue) populations at each time point. ns, p > 0.05; **,p < 0.01; ***, p < 0.001 (unpaired Student t test). **E. Diagram indicating the proportion of EdU positive progenitors after a 1 hour EdU pulse at E4.** The columns represent the percentages of EdU positive cells in electroporated progenitors (pRb/GFP double positive) in control, single CCND1, single Cdkn1c and double CCND1/Cdkn1c shRNA conditions. ns, p > 0.05; *, p < 0.05; **, p < 0.01 (unpaired Student t test). **F. Diagram indicating the percentage of PP, PN, and NN pairs of sister cells in control, single CCND1, single Cdkn1c and double CCND1/Cdkn1c shRNA conditions**. PP, PN, and NN stand for divisions producing two progenitors, one progenitor and one neuron, or two neurons, respectively. The distribution of PP, PN and NN clones between Cdkn1c *versus* control shRNA conditions was compared using the Chi-2 test, **p < 0.05; ****, p<0.005. **G. Distribution of the progenitor (pRb positive cells), neurons (HuC/D positive cells) and undefined cells (double pRb/HuC/D negative) at E4** in control, single CCND1, single Cdkn1c and double CCND1/Cdkn1c shRNA conditions. ns, p > 0.05; *, p < 0.01; ***, p<0.001 (unpaired Student t test). hae: hours after electroporation

To explore whether these changes in cell cycle parameters are responsible for the decrease in neuron production observed upon *Cdkn1c* knock-down, we targeted the CyclinD1/CDK4-6 complex, which promotes cell cycle progression and proliferation, and is inhibited by Cdkn1c. CyclinD1 loss of function by shRNA reduces the number of cycling cells in the chick embryonic neural tube (Lacomme et al., 2012; Lukaszewicz and Anderson, 2011) and favors neurogenesis in the mouse cortex, possibly through a lengthening of G1 phase (Lange et al., 2009). We hypothesized that the anti-neurogenic Cdkn1c knock-down phenotype might be caused by a failure to inhibit CyclinD1, and that a concomitant downregulation of CyclinD1 should therefore rescue, at least partially, this phenotype. Using a validated shRNA targeting chick CyclinD1 (Lukaszewicz and Anderson, 2011, Supplementary Figure 5), we compared the effect of individual and simultaneous downregulation of CyclinD1 and Cdkn1c. Remarkably, at 48 hae, whereas Cdkn1c shRNA and CyclinD1 shRNA alone respectively increased and decreased the proportion of EdU incorporation in pRb positive progenitors, the double Cdkn1c/CyclinD1 knock-down was indistinguishable from control (Figure 5E) consistent with the hypothesis that the effect of Cdkn1c on G1 duration is mediated by CyclinD1 inhibition.

We then analyzed the fate of pairs of sister cells born from progenitors dividing 24 hae in these four conditions. While CyclinD1 knock-down alone did not alter the distribution of PP, PN and NN pairs compared to the control situation, in double knock-down experiments it completely counteracted the anti-neurogenic effect of *Cdkn1c* downregulation, restoring a distribution of PP, PN and NN pairs similar to the control condition (Figure 5F). Consistently, at the population level, CyclinD1 downregulation alone did not affect the ratio of proliferating progenitors and neuron production 48 hae, but it fully rescued the *Cdkn1c* knockdown phenotype and restored the rate of neurogenesis to that of a control situation (Figure 5G).

These results demonstrate that the neurogenic activity of Cdkn1c in progenitors is largely resulting from its regulatory role on cell cycle dynamics.

## DISCUSSION

In this report, we used pseudotime reconstitution of single cell transcriptomics to identify regulators of the neurogenic transition in the chick spinal cord. After defining a set of genes clustering with the neurogenic progenitor marker Tis21, we established a “neurogenic progenitor score” that was used for differential expression analyses. We focused on one of the most significant candidates emerging from this analysis, the Cyclin/CDK inhibitor Cdkn1c. We provide evidence that *Cdkn1c* expression initiates at low levels in neurogenic progenitors before it peaks at higher levels in future neurons. Importantly, specifically altering early phase of Cdkn1c expression impairs the balance between proliferative and neurogenic modes of division. This allows us to re-interpret the role of Cdkn1c during spinal neurogenesis: while previously mostly considered as a binary regulator of cell cycle exit, we demonstrate that Cdkn1c is also an intrinsic regulator transition from the proliferative to neurogenic modes of division. This occurs through a change in the mode of division of progenitors, acting primarily via the inhibition of the CyclinD1/CDK6 complex.

Cell cycle regulators are key players of neurogenesis, controlling the ability of neural cells to either proliferate or exit the cell cycle and differentiate. Nonetheless, studies in a wide range of species have demonstrated that beyond this binary choice, cell cycle regulators also influence the neurogenic potential of progenitors, i.e the commitment of their progeny to differentiate or not (Calegari and Huttner, 2003; FUJITA, 1962; Kicheva et al., 2014; Lange et al., 2009; Lukaszewicz and Anderson, 2011a; Pilaz et al., 2009; Smith and Schoenwolf, 1987; Takahashi et al., 1995). Among cell cycle parameters, the major change observed during the neurogenic transition is a lengthening of the G1 phase: in the developing mouse cortex, progenitors positive for the neurogenic marker Tis21 display a longer G1 phase compared to Tis21 negative progenitors (Calegari et al., 2005; Calegari and Huttner, 2003; Lange et al., 2009) and apical radial glia display a shorter G1 phase than fate restricted intermediate progenitors (Arai et al., 2011). Consistently, live imaging has documented a lengthening of the G1 phase between two consecutive cycles in chick spinal neural progenitors (Molina et al., 2022). Experimental modulations of the duration of G1 phase in the dividing mother cell have revealed its pivotal role in determining the daughter cells’ capacity to self-renew or differentiate (Calegari and Huttner, 2003; Lange et al., 2009; Pilaz et al., 2009). These functional explorations have mostly focused on Cyclin/CDK complexes (Calegari et al., 2005; Calegari and Huttner, 2003; Lange et al., 2009; Pilaz et al., 2009). Intriguingly, while Cdkn1c is a key regulator of these complexes, its gain and loss of function phenotypes in the CNS have essentially been attributed to a post mitotic role in daughter cells, primarily through its function in regulating cell cycle exit (Gui et al., 2007; Mairet-Coello et al., 2012; Tury et al., 2011).

Our study combines a detailed analysis of Cdkn1c expression dynamics in progenitors and nascent neurons with functional approaches specifically targeting its activity in progenitors to resolve this apparent paradox. Using an innovative somatic knock-in strategy, we were able to tag the Cdkn1c locus with Myc epitopes and access the history of Cdkn1c transcription and translation via the Gal4-UAS reporter, and we demonstrated that the protein is expressed at low levels in neurogenic progenitors. Functional experiments dedicated to challenge specifically Cdkn1c expression in progenitors showed that reducing its expression favors a proliferative mode of division and eventually impedes neurogenesis, whereas a premature induction of low-level Cdkn1c expression is sufficient to convert proliferative into neurogenic progenitors. In addition, we showed that Cdkn1c regulates the switch to a neurogenic division mode mainly through an elongation of the G1 phase of the cell cycle resulting from the inhibition of the CyclinD/CDK6 complex (CDK4 is absent from the chicken genome).

Taken together, our study adds a new player to the complex panorama linking cell cycle dynamics and progenitor cell behavior. It reinforces the view that a faster or slower passage through the G1 phase of a progenitor is instructive for its proliferative versus neurogenic mitotic behavior, and therefore dictates the fate of its daughters. It also highlights the importance of tightly regulating the dynamics of expression of cell cycle regulators to ensure a proper neurogenic progression.

What regulates Cdkn1c expression in progenitors? Gui *et al* proposed that the strong expression of Cdkn1c that drives prospective neurons out of the cell cycle is under transcriptional control of the NeuroG2 transcription factor in the spinal cord. Indeed, a massive overexpression of NeuroG2 leads to a strong Cdkn1c upregulation and cell cycle exit (Gui et al., 2007). Our scRNAseq data indicate that NeuroG2 is already expressed at low levels in neurogenic progenitors (Figure 1D) and could therefore act upstream of Cdkn1c onset in progenitors. On the other hand, NeuroG2 protein stability and activity is controlled by the closely related p27^Kip1^/Cdkn1b (Nguyen et al., 2006). Whether Cdkn1c has a similar effect on NeuroG2 activity has not been tested directly, but it is tempting to speculate that Cdkn1c and NeuroG2 are involved in a positive feedback loop progressively leading from their moderate expression causing G1 lengthening in neurogenic progenitors to a peak of expression driving cell cycle exit in prospective neurons. Consistent with this scenario, NeuroG2 is also involved in the downregulation of cyclinD1 and cyclinE2 in spinal progenitors (Lacomme et al., 2012). Additionally, or alternatively, Cdkn1c expression may be controlled by a regulatory cascade involving the *Hes* genes. In pancreatic progenitors, Cdkn1c is transcriptionally repressed by Hes1 downstream of the Notch pathway (Georgia et al., 2006) and the same authors report a complementary expression of *Hes1* and *Cdkn1c* in the mouse neural tube. Our scRNASeq data analysis confirms a similar complementarity between Hes5 (which appears to be functionally equivalent to Hes1 in the chick spinal tube; Fior and Henrique, 2005) and Cdkn1c. The transcriptional activity of *Hes1* in pancreatic progenitors is repressed by Hes6. Interestingly, the onset of Hes6 expression shortly precedes that of Cdkn1c in our transcriptomic scRNASeq dataset. This suggests that the initiation of Cdkn1c expression in neurogenic progenitors might be triggered by the antagonistic activity of Hes6 on *Hes1/Hes5*. Further work is needed to decipher the mutual relationships regulating the dynamic expression of these factors during the neurogenic transition.

Neurogenic regulators display dynamic expression during the process of neurogenesis, and different levels in their expression may correspond to specific activities during the progression through different cellular states, as illustrated in this study with Cdkn1c. These multiple functions cannot be disentangled by complete loss-of-function or massive overexpression. Here, we circumvented these limitations via a simple and efficient somatic knock-in method that allowed us to tightly control the level and duration of Cdkn1c misexpression in a subset of progenitors in the chick embryo. The knowledge of expression dynamics of large panels of genes from single cell transcriptomics combined with our flexible method of targeted genome insertions opens the way to generalize similar customized approaches to functionally explore regulatory networks during neurogenesis. This strategy could also be extended to other contexts and animal models, enabling to bypass the lengthy process of transgenic line generation and complex crossing schemes.

One key aspect of the neurogenic transition is the reliance on asymmetric division in the early stages of neuron production. Asymmetric division of neural progenitors is an active process relying on intrinsic asymmetries in the progenitor cell involving the unequal distribution of fate determinants during mitosis (e.g. Derivery et al., 2015; Kressmann et al., 2015; Saade et al., 2017; Tozer et al., 2017). It will be important to understand whether and how Cdkn1c and other regulators of the cell cycle are involved in setting up the intrinsic polarities necessary for this process in neural progenitors. In this context, it is interesting to note that several cell cycle regulators have already been shown to play a direct role in the machinery that establishes cellular asymmetry during the division of drosophila neuroblasts (Tio et al., 2001) and sensory organ precursors (Darnat et al., 2022); on the other hand, studies in the developing mouse cortex have shown that the cyclinD2 mRNA is asymmetrically localized and inherited upon division of neural progenitors, and behaves as a fate determinant in their progeny (Tsunekawa et al., 2012). Hence, cell cycle regulators are likely to be involved at multiple levels in the process of neurogenesis, from the determination of the neurogenic competence of neural progenitors to the cellular process of asymmetric division. In this context, it will be interesting to explore whether and how Cdkn1c might control the asymmetric distribution of fate determinants that have been identified over the last years (Peyre and Morin, 2012; Saade et al., 2017; Tozer et al., 2017; Tsunekawa et al., 2012).

## MATERIALS AND METHODS

### A. Transcriptomic analysis

#### Production of scRNA-seq data

##### Sample preparation of chick cervical progenitors for single-cell RNA sequencing

Three chick embryos at 66 hours of embryonic development (E2.75, HH stage 18) were collected and dissected in ice-cold Phosphate Buffered Saline solution (PBS), transferred into ice-cold L15 medium for further dissection to retain only the cervical spinal region (spanning the length of 5 somites starting from somite number 8). To generate a single-cell suspension, dissection products were then transferred in 250µl of 37°C pre-heated papain/L15 solution (Worthington LS003126 Papain – Stock solution = 41,6mg/ml in 100ml; Working solution = 50µl of stock solution diluted in 1,5ml of L15 medium) and incubated at 37°C for 15 minutes in 2ml Eppendorf tubes. Papain was then replaced with ice-cold L15 medium, and clusters of cells were disaggregated through gentle up-and-down pipetting. Cells were then centrifuged at 300g for two minutes at 4°C. The supernatant was removed, and 500µl of new ice-cold L15 medium was added. Another round of up-and-down pipetting was performed, and cells were then sieved through 30µm filters to eliminate clumps of poorly-dissociated cells. Filtered cells were then centrifuged for four minutes at 300g, and the supernatant was replaced with 1ml of PBS containing 0.04% bovine serum albumin (BSA). Cells were then centrifuged for four minutes at 300g, 900µl of supernatant was removed and cells were then re-suspended in the remaining 100µl of solution. Quality control was assayed by counting live *versus* dead cells using Trypan blue. Samples with >90% viability were then used for the generation of scRNAseq datasets. After viability assessment, cell concentration of samples was adjusted to 1000 cells/µl.

##### Single cell transcriptomes generation, cDNA synthesis and library construction

Single-cell RNA-seq and Illumina sequencing were performed at the Ecole Normale Supérieure GenomiqueENS core facility (Paris, France). The cellular suspension (4000 cells) was loaded on 10x Chromium instrument to generate 2871 single-cell GEMs, using the manufacter’s instructions (single cell 3’ v2 protocol, 10x Genomics). Library construction was performed as per the manufacturer’s protocol and then sequenced on a NextSeq 500 device (Illumina) using paired-end (PE) 26/57, generating 533 million reads.

##### Pre-processing of chick scRNA-seq data

Primary analyses (demultiplexing, UMI processing, mapping, feature assignment and gene quantification) were performed with Eoulsan 2 (Lehmann et al., 2021). We used as references the NCBI chick reference genome assembly galGal6.fa.gz and a dedicated GTF annotation: galGal6_embryo_spinal_NT_improved.gtf. This annotation was built on top of the NCBI <u>galGal6.ncbiRefSeq.gtf.gz</u> annotation (downloaded from https://hgdownload.soe.ucsc.edu/goldenPath/galGal6/bigZips/genes/). Due to the high number of poorly annotated genes’ 3’UTRs in the chick genome, we developed a novel approach based on the re-annotation of the genome with single-cell RNA-seq data (10x Genomics short reads) and long reads bulk RNA-seq (Oxford Nanopore Technologies) from the same cell types in the chicken embryo. To this aim, we set up a genome re-annotation pipeline based on Nextflow. It takes as input the galGal6 chick reference genome and the long-reads RNA-seq data, and it outputs an improved annotation. (Lehmann et al, manuscript in preparation). This pipeline mainly relies on the use of the genome-based analysis tool IsoQuant (Prjibelski et al., 2023) for the transcript reconstruction step. We also added filtering and quality checks of the novel annotation based on the single-cell RNA seq data.

#### Biological analyses of scRNA-seq data

##### Data cleaning and preparation

The chick dataset was subjected to cleaning steps before proceeding with analyses. Filtering was performed to remove unwanted cells: cells presenting UMI counts below the 0.5th percentile and above the 99.9th percentile, more than 20% UMI counts associated with mitochondrial genes and more than 0.3% UMI counts associated with hemoglobin genes. This filtered dataset contained 2479 cells. With regard to gene filtering, we kept genes that are detected at least once in at least 3 cells. All filtering analyses were performed using Seurat v4.0.1. Scater 1.18.6.

##### Normalization and dimension reduction

Data were log-normalized with Seurat v4.0.1 function “NormalizeData”, and confounding factors such as cell cycle phases and gender were then regressed out using the function “ScaleData” (Lehmann et al., 2021). To preserve differences between proliferating and non-proliferating cells, we separated cells in two groups: “cycling” (G2/M and S) and “non-cycling” (G1/G0). Dimension reduction was then performed on scaled data, and 2D representation of the dataset (PCA and UMAP plots) were obtained. After consulting the percentage of variance explained by each dimension, we chose to keep the first 30 components.

##### Cell classification

First, using the expression of known cell population markers, we removed all cells that were neither progenitors or neurons such as cells of the mesoderm (expressing *Foxc1/2, Twist1/2, Meox1/2, Myog10*) and neural crest (expressing *Sox10*) as performed in (Delile et al., 2019). At this stage, 1878 chick cells remained. To better characterize these neural cells, we applied self-defined progenitor (P) and neuron (N) signature scores, using the Seurat function “AddModuleScore”. Scores were based on several known and newly-identified markers (originating from differential analysis performed in an initial dataset exploration). Detailed list of used marker genes is provided below.

(Progenitor genes = *Sox2, Notch1, Rrm2, Hmgb2, Cenpa, Ube2c, Hes5*; Neuron genes = *Tubb3, Stmn2, Stmn3, Nova1, Rtn1, Mapt*)

##### Clustering and differential expression

In order to identify sub-populations of cells within the population of interest, we then performed graph-based clustering using the Louvain algorithm as implemented in Seurat v4.0.1. Clustering, coupled with differential expression results (obtained using a negative binomial test) did not bring out clusters evocative of a delineation between proliferative and neurogenic progenitor populations, as cells were mainly differentiated by the patterning factors of the dorso-ventral (DV) axis. In order to find other variation sources, we designed a “denoising” strategy based on pseudotime analysis.

##### Pseudotime analysis

The pseudotime analysis was performed on the whole neural population. In order to identify genes whose expression varies over time, we relied on the “DifferentialGeneTest” function of the trajectory-inference dedicated tool Monocle3, which led to hierarchical clustering of genes along a pseudotime axis. Partitioning around medoids algorithm (PAM) was then applied to cluster cells based on similar gene expression profiles along the pseudotime. We then focused on the *Btg2/Tis21*-containing cluster to look for differentially-expressed genes (Figure 1C).

### B. Experimental Model

Fertilized eggs of JA57 chicken were purchased from EARL Morizeau (8 rue du Moulin, 28190 Dangers, France). Eggs were incubated at 38°C in a Sanyo MIR-253 incubator for the appropriate amount of time.

### Cryostat sections

For cryostat sections, chick embryos were collected at E2.25 (HH st13-14), E3 (HH st18) and E4 (HH st22) (Hamburger and Hamilton, 1992) in ice-cold PBS, then fixed over-night in 4% formaldehyde/PBS at 4°C. The following day, embryos were washed 3 times for 5 minutes in PBS at room temperature (RT). Embryos were equilibrated at 4°C in PBS/15% sucrose, then equilibrated at 42°C in PBS/15% sucrose/7,5% gelatin solution, embedded in plastic dishes containing 1ml of PBS/15% sucrose/7,5% gelatin solution and flash frozen in 100% ethanol at-50°C on dry ice, before storage at-80°C. Prior to cryostat sectioning, samples were equilibrated for 1 hour at-25°C. 20µm cryostat sections were obtained using a Leica CM3050S Cryostat and manually mounted on SuperFrost Plus microscope slides, before storage at-20°C.

### *In situ* Hybridization

For *in situ* hybridization, gelatin-mounted cryosections were first equilibrated at RT for 15 minutes, and de-gelatinized by washing slides in 37°C PBS 3 times for 5 minutes. All following steps were carried at RT unless mentioned otherwise. Slides were bathed for 20 minutes in RIPA buffer (150mM NaCl, 1% NP-40, 0.5% Na deoxycholate, 0.1% SDS, 1mM EDTA, 50mM Tris pH 8.0), post-fixed in 4% paraformaldehyde/PBS for 10 minutes, and washed with PBS 3 times for 5 minutes. Slides were then bathed in triethanolamine solution (100mM triethanolamine, acetic acid 0.25% pH 8.0) for 15 minutes and washed with PBS 3 times for 5 minutes. Subsequently, slides were pre-hybridized during one hour in 69°C pre-heated hybridization solution (50% formamide, 5X saline-sodium citrate (SSC), 5X Denhardt’s, 500 μg/ml herring sperm DNA, 250 μg/ml yeast RNA) and hybridized overnight at 69°C with the same hybridization solution in presence of the heat-denatured (95°C for 5 minutes) DIG-labelled RNA probes. The following day, slides were transferred in post-hybridization solution (50% formamide; 2x SSC; 0.1% Tween20) at 69°C for one hour, then washed in 69°C pre-heated 2x SSC solution for 30 minutes, and finally in 0.2x SSC solution at RT for 5 minutes. Slides were washed with buffer 1 (100mM maleic acid, pH 7.5, 150mM NaCl, 0.05% Tween 20) during 20 minutes at RT, blocked for 30 minutes in buffer 2 (buffer 1/10% fetal calf serum (FCS), followed by overnight incubation at 4°C with 250µl of the anti-DIG antibody (Merck #11093274910) diluted 1:2000 and other necessary primary antibodies (when additional immunostaining was needed) in buffer 2. Slides were covered with a coverslip to limit loss of solution during overnight incubation. The following day, coverslips were gently removed and slides were washed with buffer 1, 3 times for 5 minutes, and equilibrated for 30 minutes by bathing in buffer 3 (100mM Tris pH 9.5, 100mM NaCl, 50mM MgCl2). *In situ* Hybridization signal was visualized through a colour reaction by bathing slides in BM-Purple (Merck # 11442074001). The colour reaction was allowed to develop in the dark at RT during the appropriate amount of time and was stopped by bathing slides in 4% paraformaldehyde/PBS for 10 minutes. Sections were finally washed with PBS 3 times for 5 minutes before either mounting with coverslip using Aquatex or proceeding with the subsequent immunostaining protocol steps if required (see section <u>Vibratome / Cryostat sections and Immunostaining).</u>

All of RNA-probes were synthesized using a DIG RNA labelling kit (Merck #11277073910) following manufacturer’s protocol. Antisense probes were prepared from the following linearized plasmids: cHes5.1 (previously described in Baek et al., 2018b), Cdkn1c (a gift from Matthew Towers, described in Pickering et al., 2019), and cCCND1 5’ (a gift from Fabienne Pituello, previously described in Lobjois et al., 2004).

### *In ovo* electroporation

Electroporations were performed at HH13-14 by applying 5 pulses of 25V for 50ms, with 100ms in between pulses. Electroporations were performed using a square wave electroporator (Nepa Gene CUY21SC Square Wave Electroporator, or BTX ECM-830 Electro Square Porator, or Ovodyne Intracell TSS20) and a pair of 5 mm Gold plated electrodes (BTX Genetrode model 512) separated by a 4 mm interval. For bilateral electroporation of the Cdkn1c-Myc and Pax7-Cdkn1c-Myc knock-in constructs (Figure 4 and Supplementary Figure 4), the two injections were performed at 3 hours interval, the polarity of the electrode was reversed and 4 pulses of 20V were applied for each electroporation.

### Plasmids

#### RNA interference

small interfering RNA sequences against the chick version of Cdkn1c were determined using siDirect: http://sidirect2.rnai.jp/. Target sequences for Cdkn1c are as follow:

Cdkn1c shRNA 1: 5’ CGGCACCGTGCCCGCGTTCTA 3’;

Cdkn1c shRNA 2: 5’ CACGACCCGCATCACAGATTT 3’;

Cdkn1c shRNA 3: 5’ AGCGCCGTCTGCAGGAGCTTA 3’;

Cdkn1c shRNA 4: 5’ TGAGCCGGGAGAACCGCGCCG 3’;

Cdkn1c shRNA 5: 5’CGACCCGCATCACAGATTTCT 3’

Cdkn1c shRNA 6: 5’ CTCAATAAACAAAACAAAAAA 3’

Target sequences were cloned into the first hairpin of the miR30-derived structure of the pTol2-H2B-EGFP-miRNA plasmid (Peyre et al., 2011) using the following method (Das et al., 2006): 100ng of both general oligonucleotides (First hairpin primer 5’: 5’-GGCGGGGCTAGCTGGAGAAGATGCCTTCCGGAGAGGTGCTGCTGAGCG-3’ and First hairpin primer 3’: 5’-GGGTGGACGCGTAAGAGGGGAAGAAAGCTTCTAACCCCGCTATTCACCACCACTAGGCA-3’) were used together with 10ng of both target-specific oligonucleotides (Target forward sequence: 5’-GAGAGGTGCTGCTGAGCG_**FORWARDTARGETSEQUENCE_**TAGTGAAGCCACAGATGTA-3’ and Target reverse sequence: 5’-ATTCACCACCACTAGGCA_**REVERSETARGETSEQUENCE_**TACATCTGTGGCTTCACT-3’) in a one-step PCR reaction to generate a product containing the miR30 like hairpin and the chick miRNA flanking sequences. Obtained PCR products and the pTol2-H2B-EGFP-miRNA plasmid were submitted to NheI/MluI double enzymatic digestion, and purified digested products were then ligated to create CDKN1C miRNA plasmids.

The cCCND1 (chick CyclinD1) shRNA plasmid was previously described in (Lukaszewicz and Anderson, 2011), and was a kind gift of Dr Fabienne Pituello. A plasmid coding for a combination of a shRNA against Luciferase and a GFP reporter was used as a control (described in (Peyre et al., 2011). An empty pCAGGS plasmid was used to match total DNA concentrations between experimental and control electroporation mixes when needed. All miRNA and shRNA plasmids were used at 1 μg/μl except when otherwise mentioned.

#### Somatic knock-ins

Somatic knock-in of a *6xMyc-P2A-Gal4-VP16* reporter at the C-terminus of the *Cdkn1c* locus was achieved via CRISPR-Cas9-based microhomology-mediated end joining (MMEJ). The *6xMyc-P2A-Gal4-VP16* cassette in the targeting vector is flanked by 37bp 5’ and 42bp 3’ arms of homology corresponding to the genomic sequence immediately upstream and downstream of the stop codon of the CDKN1c locus. These arms of homology are flanked on both ends by a universal “uni2” gRNA target site that does not target any sequence in the chick genome (GGGAGGCGTTCGGGCCACAG; Welker et al., 2021, Petit-Vargas et al., 2024). Details of the construct and cloning steps are available upon request. The MMEJ-based knock-in method relies on the simultaneous linearization of the target locus and of the targeting vector in cells. This is achieved by coexpression of two gRNAs, one targeting the genomic locus, the other (uni2) targeting the knock-in vector. We generated a double gRNA construct that possesses two cassettes, each expressing a chimeric gRNA under control of the human U6 promoter. This vector, derived from pX330 (Cong et al., 2013); Addgene #42230), also expressed humanized spCas9 (Cas9) protein under the CBh promoter. We chose three different gRNAs located in the vicinity of the *Cdkn1c* stop codon, using the CRISPOR website (http://crispor.tefor.net/crispor.py). The sequence targeted by gRNA#1 (CTGAGCACACCCCCCGCAAG) is located 12 bases upstream of the *Cdkn1c* stop codon in the sense direction and entirely comprised in the left arm of homology. In order to avoid targeting of the knock-in vector and of the modified locus after insertion of the knock-in cassette, the target sequence for gRNA#1 was destroyed in the left arm of homology via two conservative base changes in the last base of the recognition and in the PAM (see Supplementary Figure 1). Upon initial validation of the knock-in efficiency with a UAS-nlsGFP reporter, gRNA#1 yielded the strongest GFP signal of the three gRNAs and was chosen for all subsequent experiments. A gRNA that does not target any sequence in the chick genome was used as a control (GCACTGCTACGATCTACACC; (Gandhi et al., 2017)) and did not yield any GFP signal.

Somatic knock-in of the *Cdkn1c* coding sequence in the *Pax7* locus was achieved via CRISPR-Cas9-based Homology-Directed Recombination (HDR). A *Pax7-P2A-Gal4* knock-in vector and gRNAs were first generated and validated for efficient and specific targeting at the C-terminus of the *Pax7* locus (not shown, for details see Petit-Vargas et al., 2024). In this vector, the Gal4-VP16 cassette is flanked with long left (1056bp) and right (936bp) arms of homology to the C-terminal region of *Pax7*. This vector was then modified by inserting an 846bp synthetic DNA fragment (IDT) coding for a P2A sequence, chick *Cdkn1c* and three Myc tags, immediately downstream of the Gal4-VP16 sequence. This places a *P2A-Gal4-VP16-P2A-CDKN1c-3xMyc* cassette in frame with the C-terminus of *Pax7*. The introduction of two P2A pseudo-cleavage sequences ensures that Pax7, Gal4-VP16 and CDKN1c-Myc are produced as three independent proteins from the *Pax7* locus in dorsal progenitors that have undergone homologous recombination. The *Pax7* gRNA targets the GGGCTCCTACCAGTAGAGAC sequence 16 bases upstream of the *Pax7* stop codon in the sense direction, and is entirely comprised upstream of the stop codon. In order to avoid targeting of the knock-in plasmid and re-targeting of the locus after insertion of the knock-in cassette, the gRNA target sequence was destroyed in the left arm of homology via insertion of three bases (AGA) two bases upstream of the PAM. This inserts an Arginine 5 amino acids upstream of the C-terminus of *Pax7* (see Supplementary Figure 4C). In addition to this extra amino acid, a P2A sequence is appended at the C-terminus of the Pax7 protein. We did not attempt to monitor whether this modification of the Pax7 C-terminus modifies its activity. For consistency, control experiments for the *Pax7* driven misexpression of *Cdkn1c-P2A-Gal4* were performed in a *Pax7-P2A-Gal4 knock-i*n background using the parental Pax7-P2A-Gal4 knock-in construct described in Petit-Vargas et al., 2024, which generates the same modification of the *Pax7* C-terminal sequence.

For *in ovo* knock-in experiments, the homologous recombination and gRNA vectors were each used at 0.8µg/µl. The UAS reporter plasmid (pUAS-nls-EGFP) was added to the electroporation mix at 0.3µg/µl.

### Vibratome sections

For vibratome sections, chick embryos were collected at desired stages and roughly dissected to remove membranes in ice-cold PBS, fixed for one hour in ice-cold 4% formaldehyde/PBS, and rinsed 3 times for 5 minutes in PBS at RT. Chick embryos were then finely dissected in PBS and subsequently embedded in 4% agarose (4g of agarose in 100ml of water, boiled in microwave and cooled at 50°C) until agarose became solid. Thereafter, 100 μm vibratome sections were realised using a ThermoScientific HM 650 V Microtome and collected in 6-well plates filled with cold PBS.

### Immunostaining

Sections were permeabilized in PBS-0,3% Triton for 30 minutes at RT, and then incubated with the primary antibodies diluted in the blocking solution (PBS-0,1% Triton /10% FCS) at 4°C over-night with gentle agitation. The following day, sections were washed 3 times for 5 minutes in PBS at RT, incubated 4 hours in the dark and at RT with the appropriate secondary antibodies (and DAPI if needed) diluted in PBS-0,1% Triton, washed again 3 times for 5 minutes at RT with PBS and mounted with Vectashield (with or without DAPI, depending on experiment – Vector Laboratories H-1000-10 & H-1200-10).

All immunostainings on slide-mounted cryosections were performed during and after the end of *in situ* hybridization revelation protocol. Slides were incubated with primary antibodies during the appropriate step described above in the <u>Cryostat</u> and *<u>in situ</u>* <u>hybridization</u> sections. After *in situ* hybridization revelation, slides were incubated 4 hours in the dark at RT with 250µl of appropriate secondary antibodies (and DAPI, if needed) diluted in PBS-0,1% Triton, washed again 3 times for 5 minutes at RT with PBS and mounted with Aquatex.

Primary antibodies used are: chick anti-GFP (GFP-1020 – 1:2000) from Aves Labs; goat anti-Sox2 (clone Y-17 – 1:1000) from Santa Cruz; rabbit anti-pRb (Ser807/811 – 1:1000) from Cell Signaling; mouse anti c-myc tag (Clone 9E10 - 1:100) from Sigma-Aldrich; rabbit anti DsRed (Polyclonal - 1: 400) from Takara Bio; mouse anti-HuC/D (clone 16A11 – 1:50) from Life Technologies. Secondary antibodies coupled to Alexa Fluor 488, Cy3 or Alexa Fluor 649 were all obtained from Jackson laboratories and all used at a 1:500 dilution.

### EdU labelling

Proliferating progenitors in the neural tube were labelled with 5-ethynyl-2′-deoxyuridine (EdU) via *in ovo* incorporation. Before EdU injection, membranes surrounding the embryos were slightly opened using forceps. For one-hour pulse experiments, 100μl of a 500 μM solution of EdU diluted in PBS was deposited directly on the embryo through the opened membranes. For cumulative EdU labelling, embryos were incubated with EdU for the appropriate amount of time before collection. In this context, 100μl of a 500 μM solution of EdU diluted in PBS was deposited every 6 hours through the previously opened space after initial injection. After collection, embryos were subsequently processed following the vibratome sections protocol. Revelation of EdU incorporated in progenitors was carried out on vibratome sections after the permeabilization step, using the Click-iT EdU imaging kit (Invitrogen) according to manufacturer’s protocol.

### FlashTag preparation and injection

A 1mM stock solution of CellTrace Far Red (Life Technologies, #<u>C34564-</u> (Baek et al., 2018)) was prepared by adding 20μl of DMSO to a CellTrace Far red dye stock vial. A working solution of 100μM was subsequently prepared by diluting 1μl of stock solution in 9μl of 37°C pre-heated PBS, and injected directly into E3 chick neural tubes. The eggs were resealed with parafilm and embryos were incubated at 38 °C for the appropriate time until dissection.

### Image Acquisition

Transverse sections of chick embryo neural tubes after *in situ* hybridization and/or immunofluorescence were obtained either on a confocal microscope (model SP5; Leica) using 40× and 63× (Plan Neofluar NA 1.3 oil immersion) objectives and Leica LAS software, or on an inverted microscope (Nikon TiEclipse) equipped with a Yokogawa CSU-WI spinning disk confocal head, a Borealis system (Andor Technologies) and an sCMOS Camera (Orca Flash4LT, Hamamatsu) using a 40× objective (CFI Plan APO LBDA, NA 0.45, Nikon) or a 100× oil immersion objective (APO VC, NA 1.4, Nikon) and micromanager software (Edelstein et al., 2010). For image processing, data analysis and quantifications, we used the Fiji software to adjust brightness and contrast.

### Image Quantifications

In the ventral motor neuron domain of the neural tube, progenitors differentiate earlier than in any other region of the neural tube. Thus, to reason on a more homogeneous progenitor population, we restricted all our analysis to the dorsal one half or two thirds of the neural tube. All cell counting in this study were performed manually.

### Status of proliferation/differentiation balance at tissue level

HH13-14 chick embryos were electroporated with gene-specific or control shRNAs co-expressing a Histone-GFP reporter, and status of the proliferation/differentiation balance in the GFP positive electroporated population was analysed one or two days after electroporation.

Unambiguous identification of cycling progenitors and postmitotic neurons is notoriously difficult in the chick spinal cord, as there are no reliable reagents that label these populations: markers of neurons, such as HuC/D of bIII-tubulin (TujI) are not detected during the first hours of neural differentiation; on the other hand, markers of progenitors usually either do not label all the phases of the cell cycle (eg. Phospho-Rb, thereafter pRb), or persist transiently in newborn neurons (eg. Sox2). With these limitations in mind, we used antibodies against HuC/D to label neurons and phosphorylated Rb to identify the progenitor population. This leaves a population of “undetermined cells” that are negative for both markers, and that can be either progenitors in G1 phase before the restriction point and are therefore still pRb negative, or newborn neurons that have not yet activated HuC/D. For conditions analysed using a combination of GFP, pRb and HuC/D primary antibodies, three ratios were determined. Progenitor ratio was obtained by dividing the number of shRNA electroporated (GFP positive) and pRb positive/HuC/D negative cells by the total number of electroporated cells (GFP positive). Neuron ratio was obtained by counting the number of shRNA electroporated (GFP)

HuC/D positive cells over the total number of electroporated cells (GFP positive). The ratio of “undetermined cells” was obtained by dividing the number of shRNA electroporated (GFP positive) cells that were negative for both pRb and HuC/D by the total number of electroporated cells (GFP positive).

### Cumulative EdU incorporation in *Cdkn1c* knock-down and control progenitors

Chick embryos were electroporated at E2 with either control or Cdkn1c shRNA vectors and EdU injections were performed *in ovo* starting at E3 and then every 6 hours to cover the whole cell cycle.

At each measured timepoint (1h, 4h, 7h, 10h, 12h, 14 and 17h after the first EdU injection), we quantified the number of EdU positive electroporated progenitors (triple positive for EdU, pRb and GFP) over the total population of electroporated progenitor cells (pRb and GFP positive) (Figure 3B). The average values of cycling progenitors obtained for each time point were then plotted to construct the graphs. The numbers of embryos, sections and cells quantified for each timepoint in each condition is detailed below.

Cdkn1c shRNA condition: A minimum of 730 cells, collected from 3 to 5 embryos were analysed for each timepoint, from 1 to 2 different experiments.

Control shRNA condition: A minimum of 689 cells, collected from 2 to 5 embryos were analysed for each timepoint, from 1 to 2 different experiments.

### Quantification of progenitors in S phase at a given time point (1 hour EdU pulse)

Control shRNA, Cdkn1c shRNA, cCCND1 shRNA and a reporter of electroporation (GFP), and a combination of Cdkn1c and cCCND1 shRNAs were electroporated in HH13-14 chick embryos. Two days after shRNAs electroporation, we injected EdU *in ovo* one hour before collecting the embryos. We then labelled transverse sections for EdU incorporation as described in EdU labelling section. We quantified the proportion of progenitors in S phase in the shRNA conditions (GFP/EdU/pRb positive cells) over the global population of electroporated progenitors (GFP/pRb positive). For each condition, a minimum of 1594 cells collected from 3 to 6 embryos were analysed. Post-EP = post-electroporation.

### G1 analysis of neural progenitors at the cell level

We used the FlashTag (FT) technique, based on the ability of the cell-permeant dye CellTrace Far Red (Life Technologies, #C34564) to fluorescently label intracellular proteins. Previous experiments in the embryonic chick have shown that upon direct injection in the neural tube, FT dyes preferentially enter progenitor cells undergoing mitosis near the apical surface and that this incorporation only occurs during a 15-30 minutes’ time window (Baek et al., 2018). Since FT fluorescence is preserved in daughter cells after mitosis, this dye offers a convenient means to synchronously label a cohort of cells dividing at the time of injection and follow their progeny.

Using a combination of FT injection and cumulative EdU incorporation allows to monitor precisely the average length of the G1 phase. Daughter cells from FT positive progenitors enter G0/G1 phase just after mitosis, and will start incorporating EdU only when entering S phase. For each cell in a FT cohort, the time window between FT injection and the beginning of EdU incorporation corresponds to the duration of the G1 phase. One day after electroporation, FT and EdU were injected simultaneously and embryos were collected at different time points after injection to identify the time at which all cells in the FT cohort have exited G1 and entered S phase. For time points over 6 hours, an additional EdU injection was performed after 6 hours to cumulatively label the whole population of cycling cells.

We quantified the number of electroporated (GFP positive) progenitors (pRb positive) having incorporated EdU and FT (GFP/pRb/EdU/FT quadruple positive) relative to the number of FT electroporated progenitors (GFP/pRb/FT triple positive). At each time point, the percentage of quadruple positive cells represents the proportion of progenitors having completed their G1 phase. This percentage reaches a plateau when all the progenitors in the FT cohort have entered S phase. Therefore, an experimental FT cohort that reaches the plateau faster than the control FT cohort has a shorter G1 phase duration.

### Clonal analysis of sister cell identities and mode of division

We first determined the time point after mitosis at which pRb becomes a reliable progenitor marker by monitoring the time window after which all progenitors in a synchronized cohort of cells undergoing mitosis reach the restriction point/late G1 stage, as determined by pRb immunoreactivity. Using FT to label a cohort of pairs of sister cells that perform their division synchronously at E3, we counted the proportion of pairs in the cohort that contained 0, 1 or 2 cells positive for pRb at different time points after FT injection. The distribution between these three categories should reach a plateau when all the progenitors in the cohort have passed the restriction point and have become positive for pRb. At E3, this plateau was reached between 4h30 and 6 hours after FT injection, and the distribution of pRb immunoreactivity within pairs of FT positive sister cells was stable at later time points, indicating that from 6 hours after FT injection, the proportions of FT pairs with 0, 1, or 2 pRb positive cells respectively correspond to the proportions of NN, PN and PP pairs in the cohort. We therefore choose to perform clonal analysis in embryos harvested 6 hours after FT injection.

Chick embryos were electroporated at E2 with the relevant shRNAs. One day later FT was injected in the neural tubes in order to follow the progeny of isochronic dividing neural progenitors. Cell identity of transfected GFP positive cells was determined as follows: cells positive for pRb and FT were classified as progenitors and cells positive for FT and negative for pRb as neurons. In addition, a similar intensity of both the GFP and FT signals within pairs of cells, as well as their proximity within the tissue were used as criteria to further ascertain sisterhood. Using these criteria to identify pairs of sister cells, the mode of division used by their mother cell was determined as follows: symmetric proliferative if the two daughter cells were attributed the progenitor identity (PP); asymmetric if one of the daughters was a progenitor and the other daughter a neuron (PN); and terminal neurogenic if the two daughter cells had a neuronal identity (NN).

### Statistical analyses

The number of embryos and analysed cells or sections are indicated above. All data processing and statistical analyses were performed using Excel and GraphPad Prism software and are indicated in Legends to Figures.

### Contact for Reagent and Resource Sharing

As Lead Contact, Xavier Morin (Institut de Biologie de l’Ecole Normale Supérieure) is responsible for all reagent and resource requests. Please contact Xavier Morin at with requests and enquiries.

#### Key Resources Table – STAR METHODS

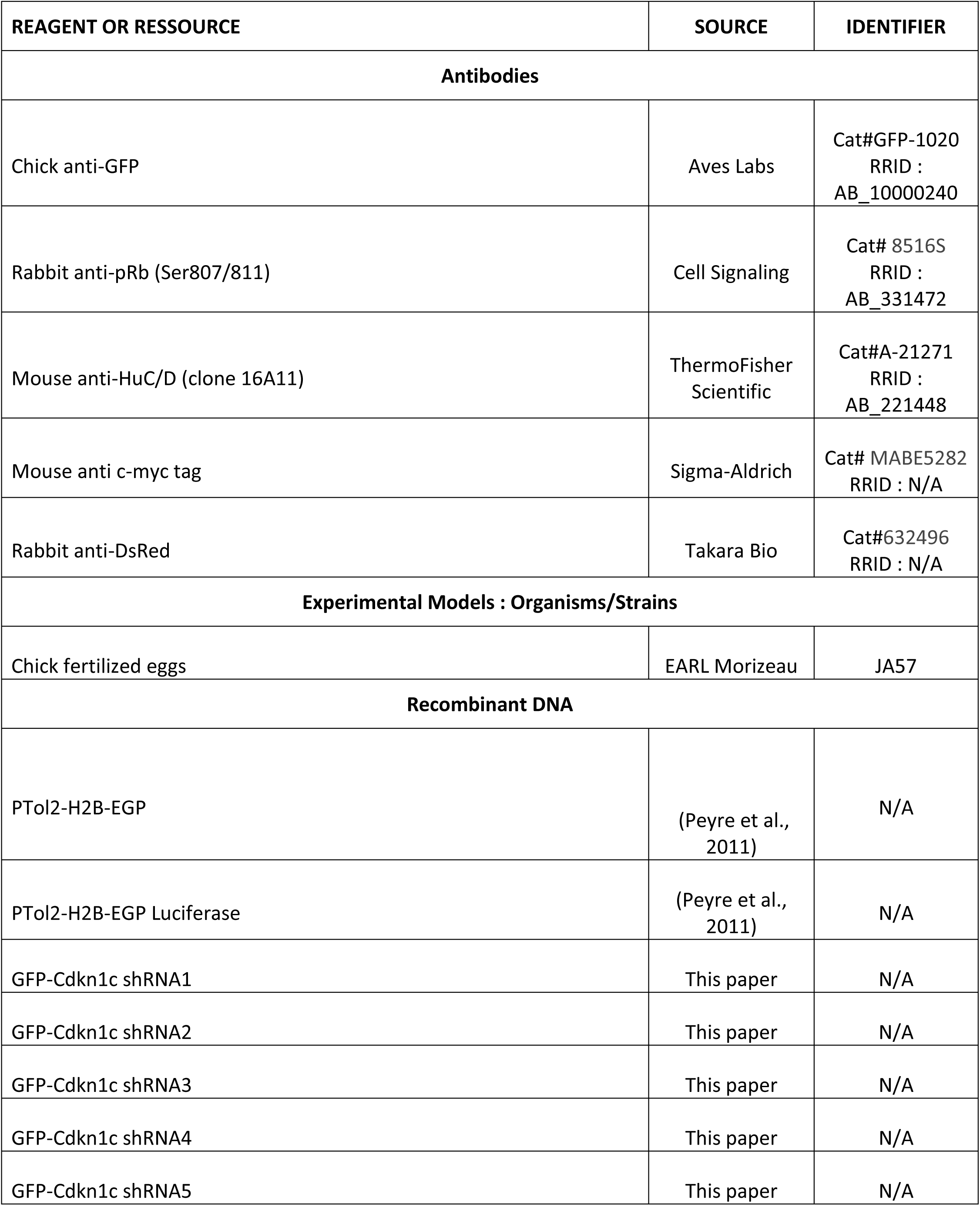
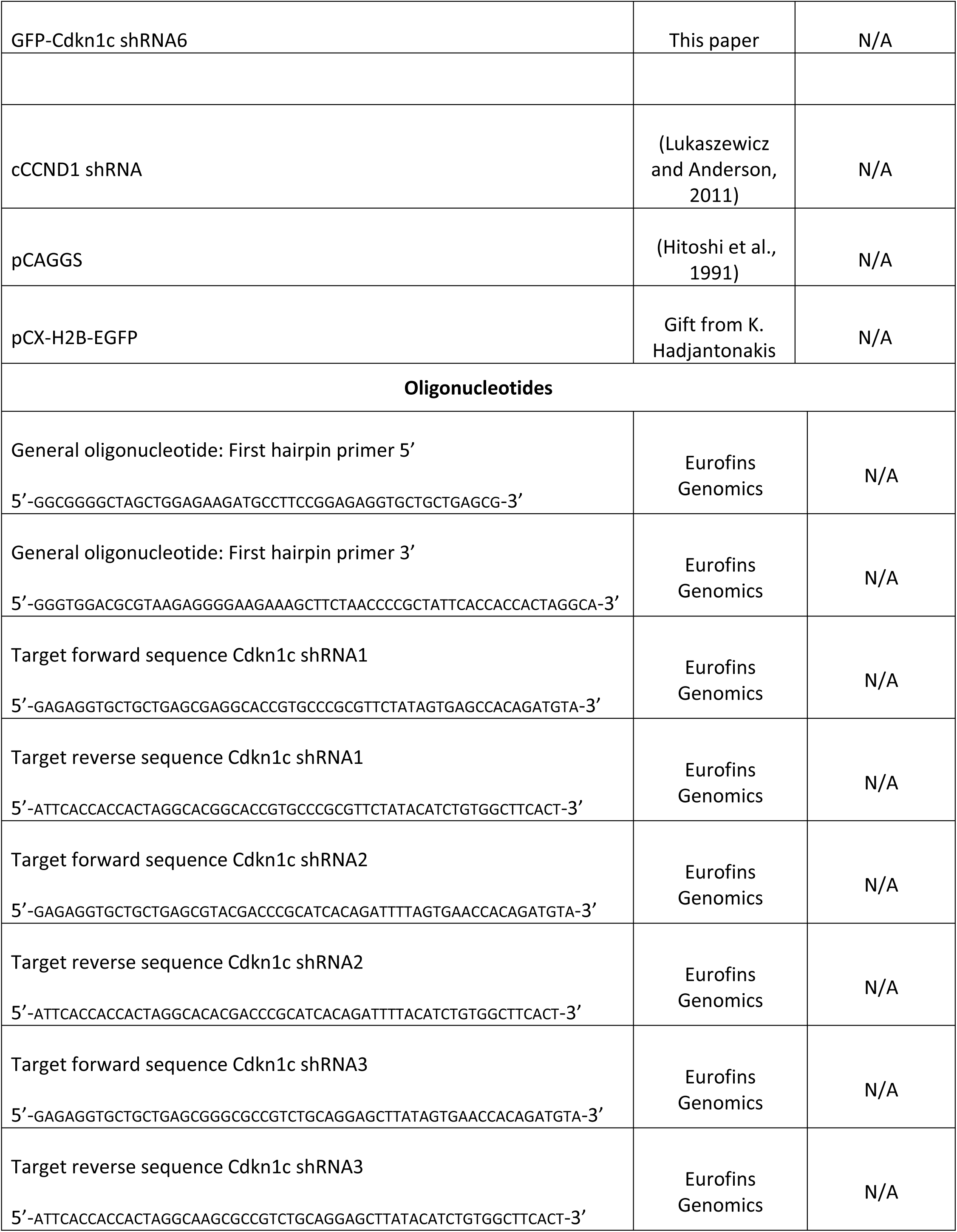
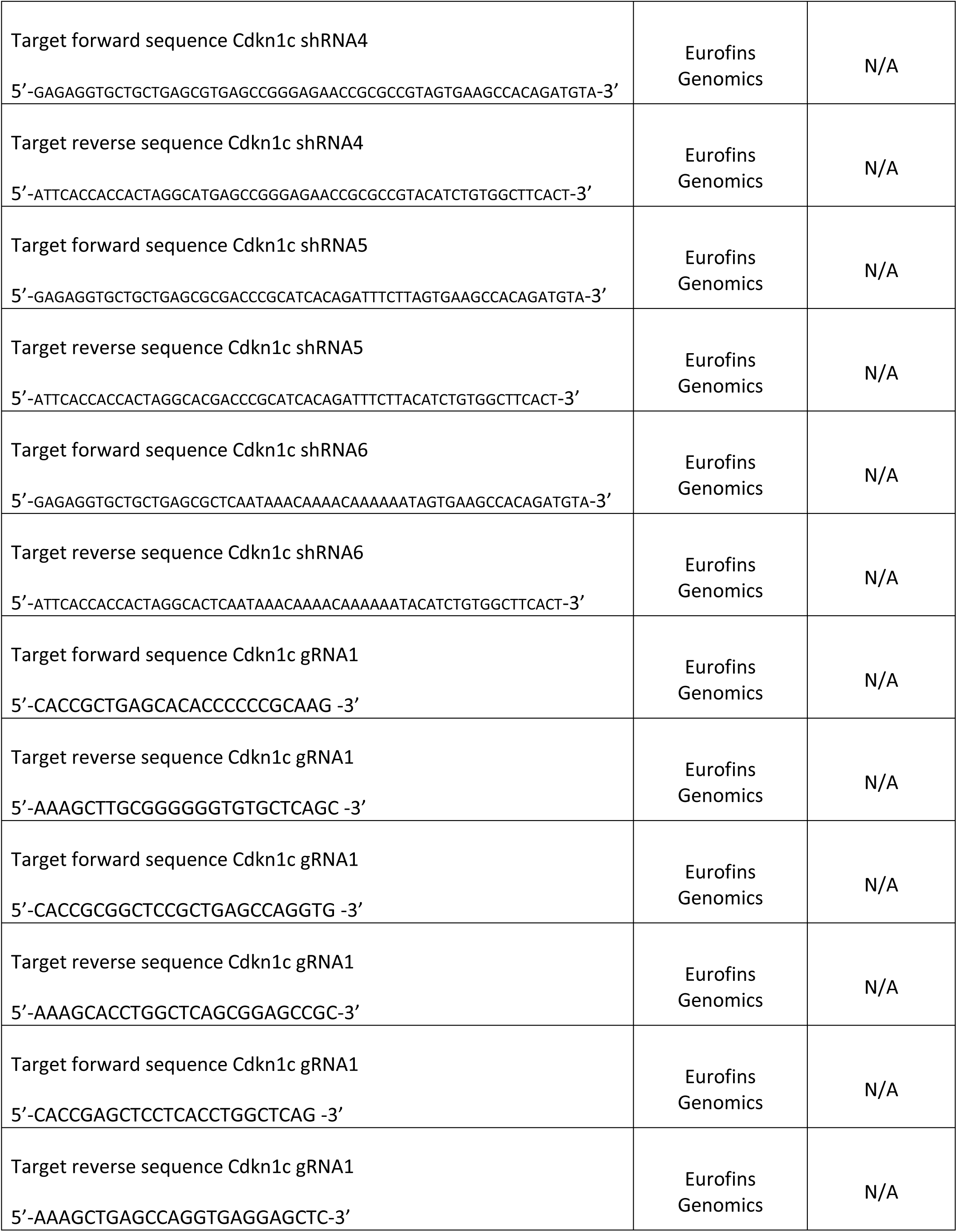
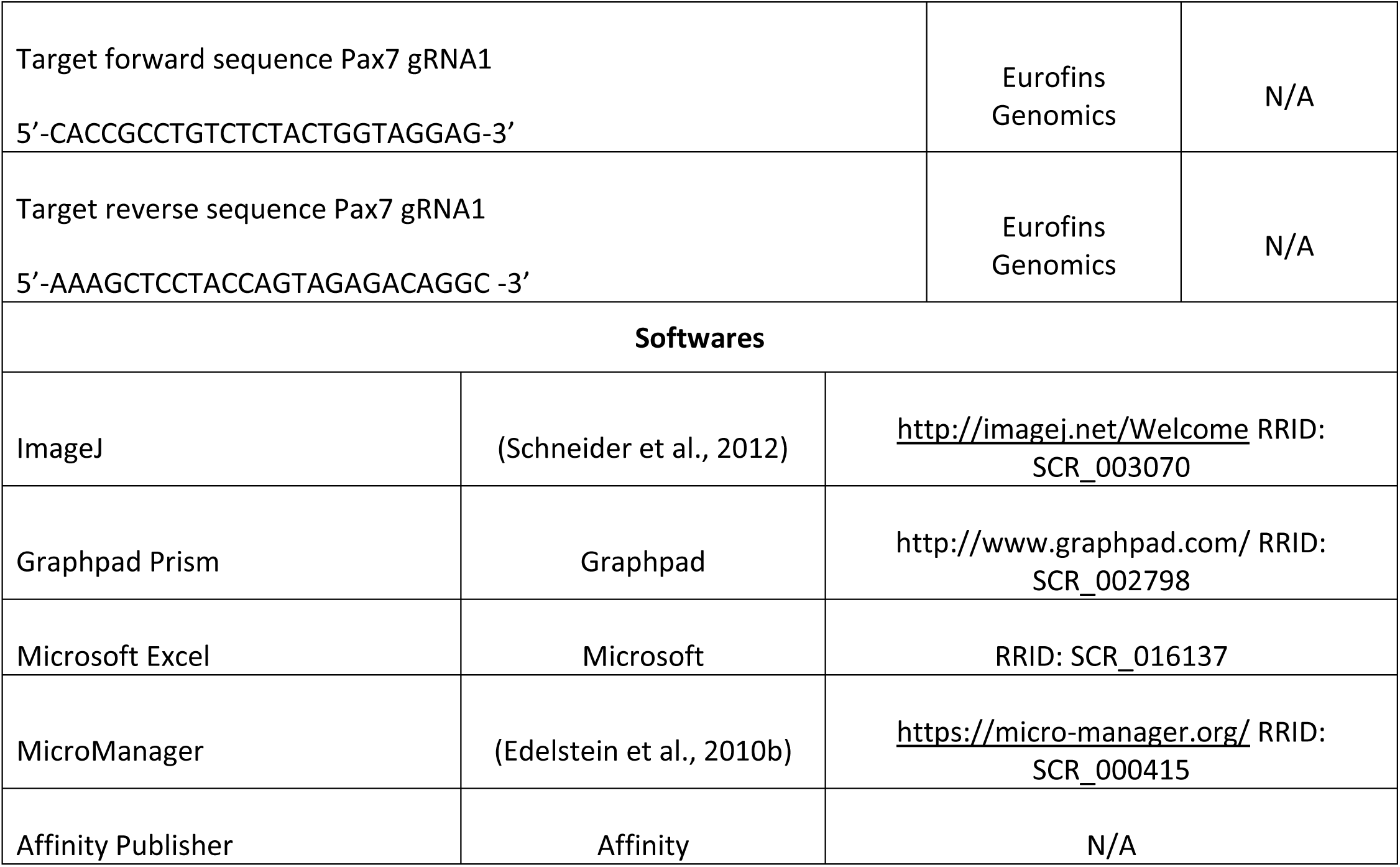

## ACKNOWLEDGEMENTS

We thank our colleague Samuel Tozer for discussions and critical reading of the manuscript. We thank Fabienne Pituello and Matthew Towers for plasmids. We thank Benjamin Bunel for the original drafting of the figures. This work was supported by grants from the Fondation pour la Recherche Medicale (FRM EQU202003010547), the Labex MEMO LIFE, the Fondation Cino del Duca to XM, the Institut Universitaire de France to MTC and a joint grant from the Agence Nationale pour la Recherche (SYMASYM ANR-18-CE16-0021-01) to MTC and XM. BM was supported by doctoral grants from the French Ministry of Higher Education and Research (MESR) and the Labex MEMO LIFE. This work has received support under the program « Investissements d’Avenir » launched by the French Government and implemented by the ANR, with the reference ANR-10-LABX-54 MEMO LIFE. The GenomiqueENS core facility was supported by the France Génomique national infrastructure, funded as part of the “Investissements d’Avenir” program managed by the Agence Nationale de la Recherche (contract ANR-10-INBS-09). The authors declare no competing financial interests.

## AUTHOR CONTRIBUTIONS

Conceptualization – EF, XM

Data curation – NL

Formal analysis – NL, BM

Funding acquisition – MT-C, XM

Investigation – BM, FC, RG, KB

Methodology – NL, MT-C, EF, XM

Project administration – MT-C, EF, XM

Software – NL

Supervision – MT-C, EF, XM

Validation – EF, XM

Visualization – BM, NL, EF, XM

Writing – original draft – EF, XM

Writing – review & editing – NL, MT-C, EF, XM

---

**Supplementary Figure 1:**
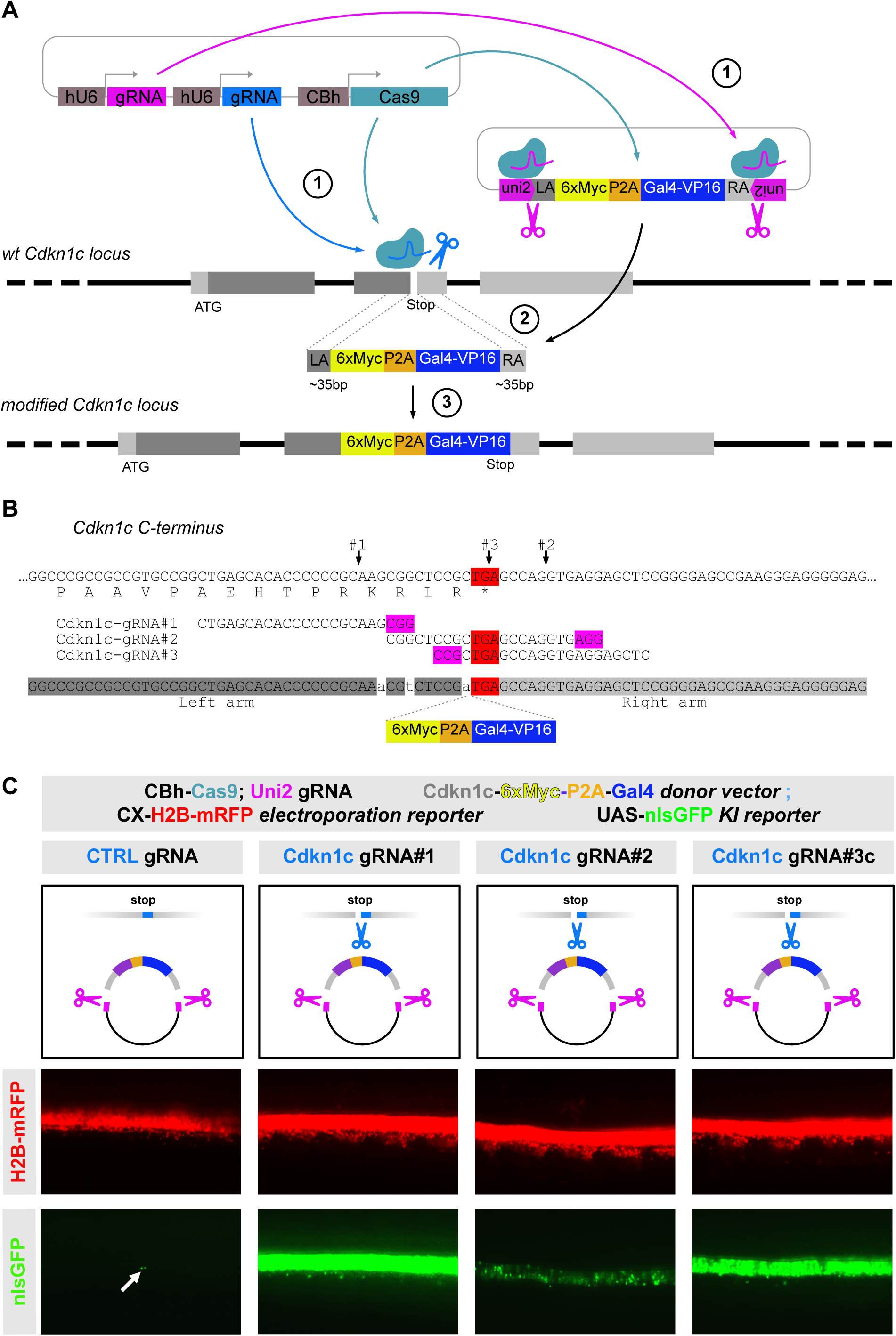
Somatic knock-in strategy to target the endogenous *Cdkn1c* locus with Myc tags and monitor the dynamic expression of the protein in spinal cord progenitors. **A. “micro-homology mediated end joining” (MMEJ) strategy used for the somatic knock-in.** The donor plasmid carries the short arms of homology (<35bp) to the *Cdkn1c* locus at the level of the C-terminus, flanking a sequence consisting of 6 Myc tags in frame with *Cdkn1c* coding sequence, a P2A pseudo cleavage sequence and the Gal4-VP16 synthetic transcription factor. This donor cassette is flanked on both sides by target sites for a “universal” guide RNA (uni2 gRNA) designed to trigger linearization of the vector and to release the donor cassette as a linear double-stranded DNA fragment in the electroporated cells. Somatic knock-in is achieved by the coelectroporation of a CRISPR/Cas9 vector that expresses the Cas9 protein, the uni2 gRNA targeting the donor vector for linearization, and the gRNA targeting the *Cdkn1c* locus. **B. Details of the targeted genomic sequence for Myc tagging of the endogenous** *Cdkn1c* **locus.** Genomic sequence at the level of *Cdkn1c* C-terminus (top), sequence of the three gRNAs (middle), and sequence of the arms of homology used in the MMEJ construct (bottom, sequence highlighted in blue). The three bases highlighted in white represent silent base changes introduced in the left arm of homology to prevent recognition and cleavage of the donor vector by gRNA#1. Arrows indicate the theoretical cut sites of the three gRNAs on the target locus. **C. Validation of the efficiency and specificity of the knock-in strategy:** the donor vector was coelectroporated with CRISPR/Cas9 vectors expressing either a control gRNA (CTRL gRNA) or one of the three gRNAs targeting the *Cdkn1c* locus. A UAS-nlsGFP vector was included in the electroporation mix to report expression of the Gal4-VP16 transcription factor in addition to an electroporation reporter (CX-H2B-mRFP). One representative embryo is shown for each condition, with similar electroporation level (red). gRNA#1 led to a strong GFP signal (green), showing the greatest efficiency. gRNA#3 was slightly less efficient, and gRNA#2 yielded a much lower signal. Specificity is demonstrated by the virtual absence of background GFP signal when the control gRNA is used (white arrow points to two GFP positive cells observed in the control embryo).

**Supplementary Figure 2:**
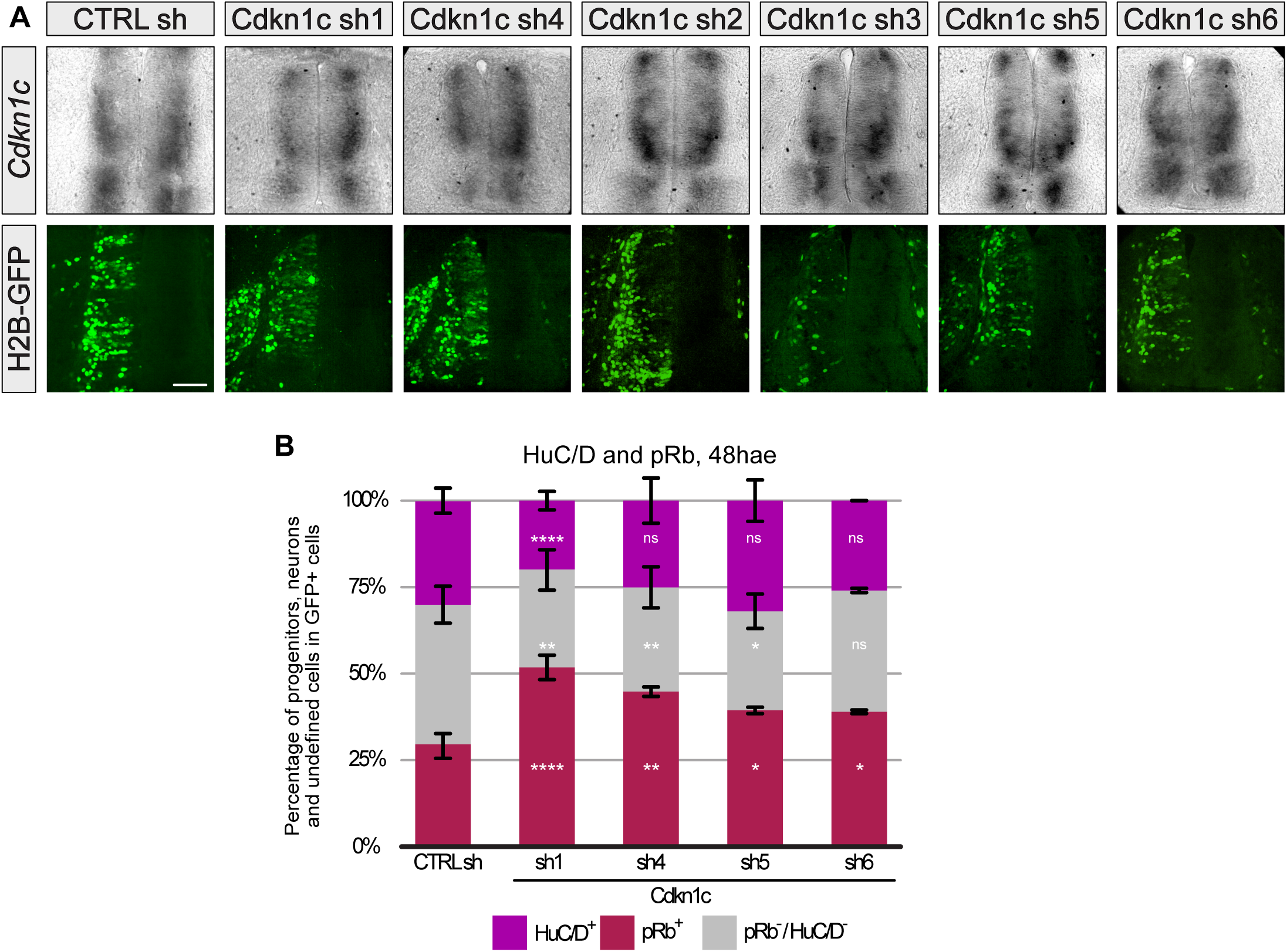
Partial knock down of Cdkn1c expression in spinal neural progenitors and prospective neurons via shRNA delays neurogenesis **A. mRNA expression of Cdkn1c in chick embryonic neural tube after electroporation of each of the six shRNAs**. *In situ* hybridization and GFP immunofluorescence on the same cryosection of thoracic region of chick embryo. Upper panel: visible downregulation of *Cdkn1c* mRNA was only observed with shRNA1 and to a lesser extent shRNA4 conditions, while comparable mRNA expression to the control condition was observed with the other shRNAs (compare left versus right hemitube). Lower panel: Corresponding level of electroporation for each embryo (GFP immunofluorescence). Scale bar: 50µm **B. Distribution of the pRb positive progenitors (red), HuC/D positive neurons (green) and undefined cells (double pRb/ HuC/D negative, gray) in shRNAs 1, 4, 5, 6 or control conditions at E4, 48 hours after electroporation (hae)**. (values for CTRL and shRNA1 are identical to Figure 2B). ns, p > 0.05; *, p < 0.05; **, p < 0.01; ****, p < 0.001 (unpaired Student’s t test relative to CTRL shRNA).

**Supplementary Figure 3:**
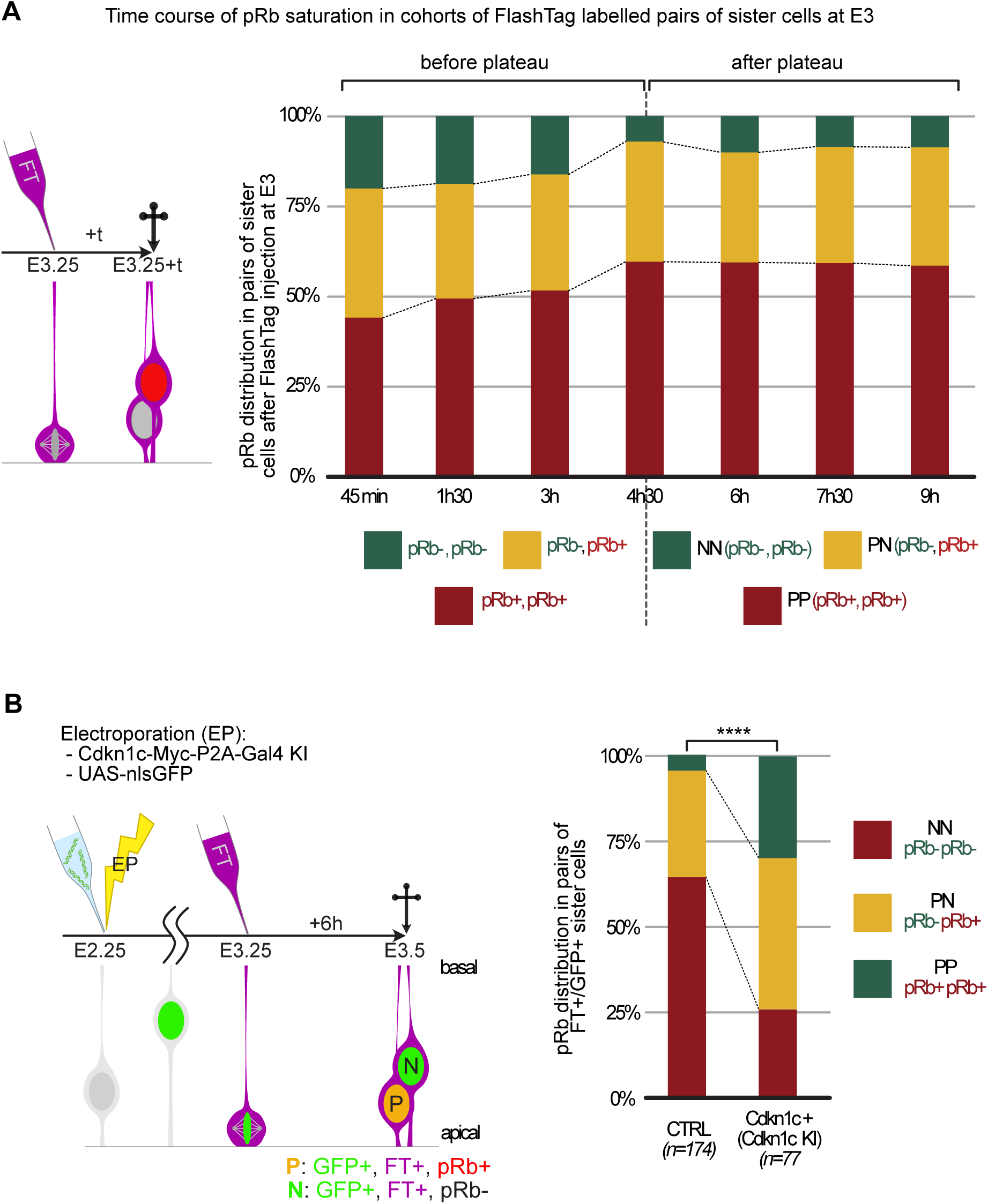
pRb is a reliable marker of progenitor cells six hours after mitosis **A. Left: Experimental scheme for the determination of Rb phosphorylation status in pairs of sister cells.** Wild type E3.25 embryos are injected with FlashTag (FT). They are collected at different timepoints (t) after injection to determine at which timepoint pRb positivity in sister pairs reaches a plateau. **Right: Time course of pRb expression in pairs of sister cells at consecutive time points after FlashTag injection.** FlashTag injection was performed at HH17-18 (E3.25) to label a synchronous cohort of mitotic progenitors and embryos were harvested at the indicated timepoints after injection. Thoracic vibratome sections were immunostained with anti-pRb antibody to evaluate the pRb status in pairs of FlashTag positive sister cells. The proportion of pairs with two pRb positive cells (green), one pRb positive cell (yellow) or zero pRb positive cell (red) reaches a plateau after 4h30, indicating that after that time point, pRb positivity becomes a reliable marker of progenitor status in FlashTag labelled pairs of sister cells. **B. Cdkn1c positive progenitors are more neurogenic than the overall progenitor population**. Left: Experimental scheme for the clonal analysis. A knock-in of the Gal4 reporter in the *Cdkn1c* locus was performed at E2.25 and FlashTag (FT) was injected 24 hours later. Sister cells born from Cdkn1c-positive progenitors dividing at the time of FlashTag injection were identified on the basis of the expression of a UAS-nlsGFP reporter and FlashTag positivity. The distribution of of PP, PN, and NN pairs is significantly different between the *Cdkn1c* knock-in progenitors and the contralateral side of the same transverse sections, indicating that the Cdkn1c positive population of progenitors is significantly more neurogenic than the whole population at that stage. Chi-2 test, ****, p<0.005.

**Supplementary Figure 4:**
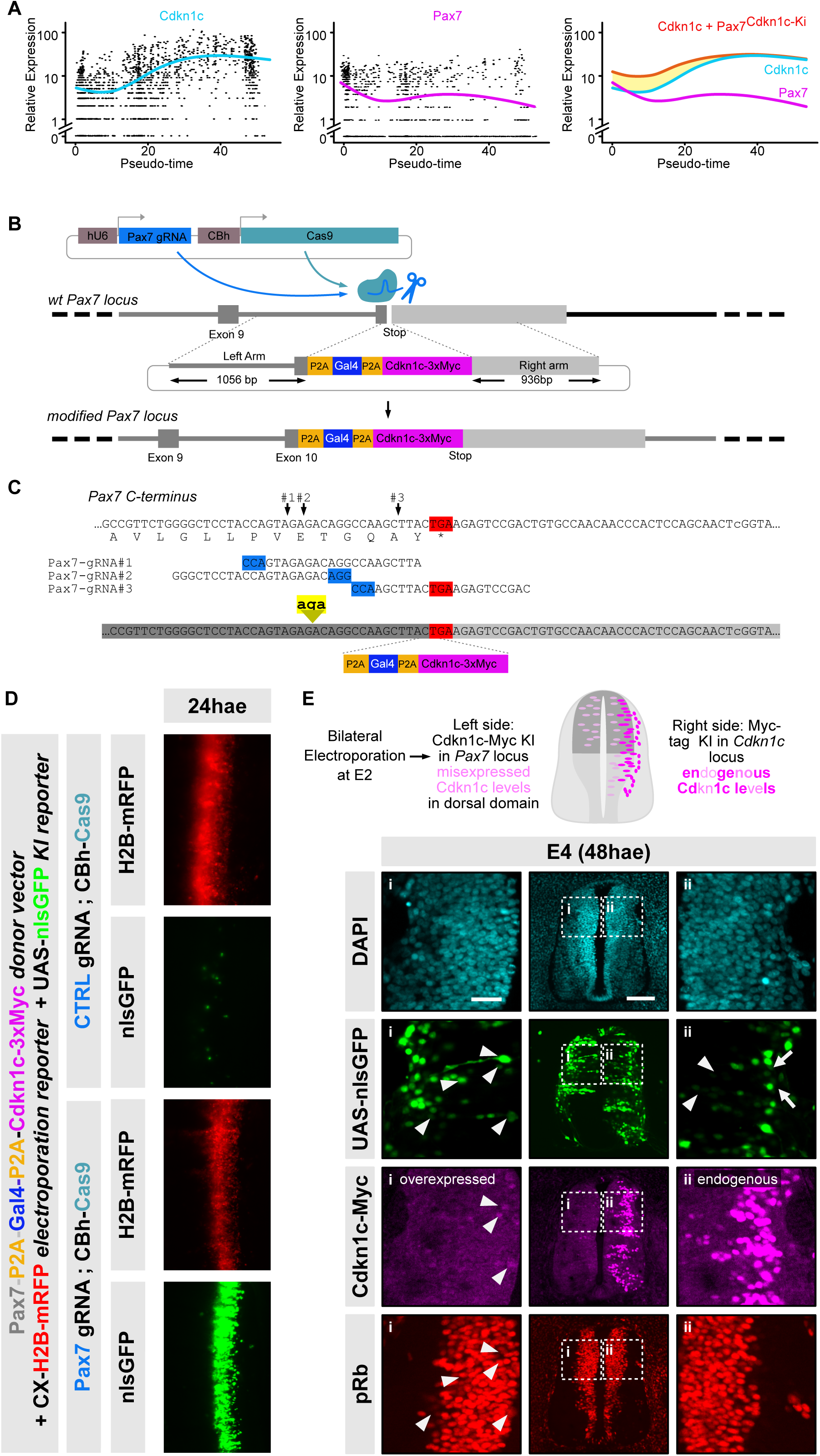
*Cdkn1c* knock-in at the *Pax7* locus leads to a premature low-level expression of Cdkn1c in dorsal neural progenitors. **A. Expression levels of *Cdkn1c* and *Pax7* transcripts along the pseudo-time axis from the chick scRNASeq analysis**. *Cdkn1c* expression has very low levels in “early” progenitors (left part of the pseudotime axis) and increases in more mature progenitors, before peaking in differentiating neurons. Pax 7 expression levels in “early” progenitors are slightly higher than those of *Cdkn1c*. The right panel shows an estimated cumulative expression (red) of endogenous (blue) and *Pax7*-driven (magenta) *Cdkn1c* levels upon knock-in of *Cdkn1c* coding sequences in the *Pax7* locus, which should result in premature expression in early progenitors, but no overexpression in newborn neurons. The scale for “Relative Expression” is logarithmic **B. Principle of the “homology directed repair” (HDR) strategy used to drive low-level misexpression of *Cdkn1c*-Myc and Gal4-VP16 in dorsal progenitors from the *Pax7* locus.** Somatic knock-in is based on a donor plasmid which carries long arms of homology (∼1Kb) to the *Pax7* locus at the level of the C-terminus in exon 10. The arms of homology flank an “in-frame” knock-in cassette that consists in a P2A pseudo-cleavage site, the Gal4-VP16 synthetic transcription factor, a second P2A pseudo-cleavage site and the *Cdkn1c* coding sequence fused to three C-terminal Myc tags. This knock-in approach requires in addition the electroporation of a second vector expressing the Cas9 protein and a gRNA that targets the genomic region of *Pax7* upstream of the stop codon. Upon successful knock-in insertion, Pax7 (gray), Gal4-VP16 (blue) and Cdkn1c-Myc (magenta) coding sequences will be transcribed from the *Pax7* locus and co-translated. The insertion of P2A pseudo-cleavage sites (orange) between the three sequences will ensure that all three proteins are present as independent proteins. **C. Details of the targeted genomic sequence at the C-terminus of the *Pax7* locus.** Genomic sequence at the level of *Pax7* C-terminus (top), sequence of the three independent gRNAs targeting this region (middle), and sequence of the arms of homology surrounding the knock-in cassette (bottom). To avoid possible targeting of the donor arms by gRNAs#1 and #2, three bases (highlighted in yellow) were inserted 5 amino acids upstream of the *Pax7* stop codon. Note that this introduces an Arginine residue (AGA) in the *Pax7* sequence. **D. Validation of the efficiency and specificity of the knock-in strategy.** Imaging of the neural tube directly *in ovo*. The donor vector was coelectroporated together with a dual vector expressing the Cas9 nuclease and either a control gRNA (CTRL gRNA) or the gRNA targeting the *Pax7* locus. A UAS-nlsGFP vector was included in the electroporation mix to report expression of the Gal4-VP16 transcription factor. Finally, an electroporation reporter (CX-H2B-mRFP) was added to monitor the quality of electroporation. One representative embryo is shown for each condition, with similar electroporation level (red). Specificity is demonstrated by the virtual absence of background GFP signal when the control gRNA is used (compared to the massive GFP signal with *Pax7* gRNA, only few GFP positive cells are observed in the control embryo). In addition, specificity was demonstrated by the dorsally restricted expression of the GFP signal in the Pax7 domain on transverse section (see for example, GFP signal in the right hemitube in panel E). **E. *Pax7*-driven exogenous expression of Cdkn1c in dorsal progenitors mimics the levels of Cdkn1c expression in neurogenic progenitors at E4.** A bilateral electroporation scheme was used to compare *Pax7*-driven levels of Cdkn1c expression (electroporation 1, right side hemi-tube, knock-in of Cdkn1c-Myc in the *Pax7* locus) with endogenous Cdkn1c levels (electroporation 2, left side hemi-tube, knock-in of a Myc tag in the *Cdkn1c* locus). The level of Cdkn1c-Myc (magenta) expression driven by *Pax7* is low and restricted to the ventricular region, where it is comparable to the endogenous levels of Cdkn1c-Myc expression in the contralateral side (i and ii, arrowheads). Note that although very few cells with a detectable Cdkn1c-Myc expression are observed in the misexpressed condition, the UAS-nlsGFP reporter is widely expressed, indicating strong electroporation and knock-in efficiency. This strong GFP signal, compared to the weak Myc signal, is explained by a differential stability and posttranslational regulation between Cdkn1c-Myc and Gal4-VP16, and by amplification of the GFP fluorescence via the Gal4/UAS system. Scale bars, 100µm in central column and 30 µm in close ups.

**Supplementary Figure 5:**
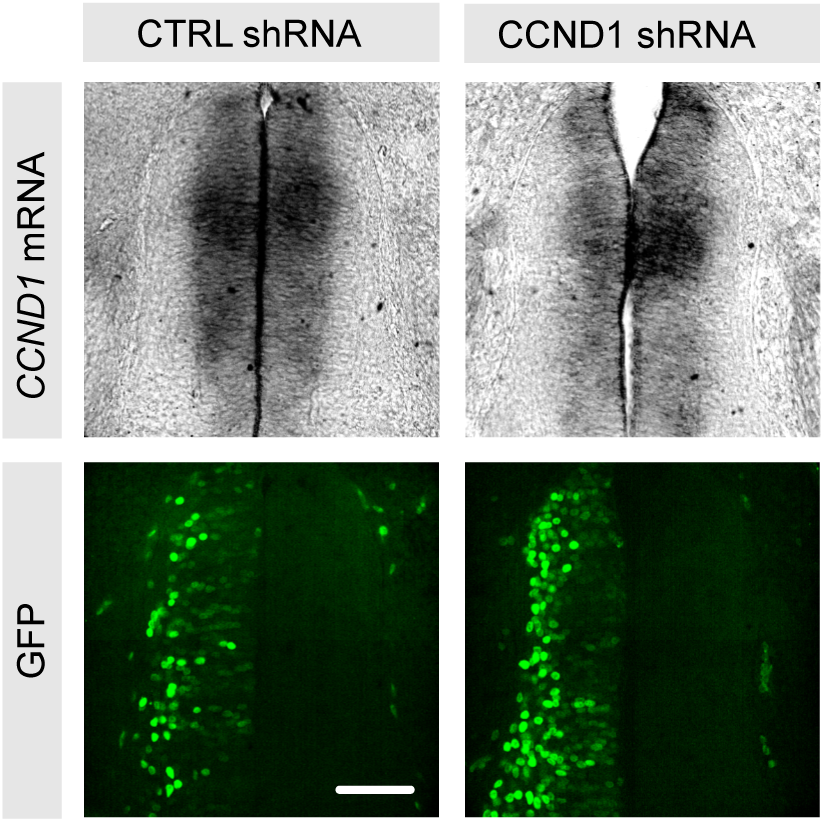
Decreased *CCND1* mRNA expression in chick embryonic neural tube one day after CCND1 shRNA electroporation. *In situ* hybridization of a *CCND1* antisense probe on transverse cryo-sections of chick embryonic neural tube followed by an anti-GFP immunostaining to reveal electroporated cells. Scale bars: 100µm

